# Real-time genomic pathogen, resistance, and host range characterization from passive water sampling of wetland ecosystems

**DOI:** 10.1101/2025.09.05.674394

**Authors:** Albert Perlas, Tim Reska, Alberto Sánchez-Cano, Cristina Mejías-Molina, Daniel Gygax, Sandra Martínez-Puchol, Marta Rusiñol, Elias Eger, Katharina Schaufler, Ursula Höfle, Guillaume Croville, Guillaume Le Loc’h, Jean-Luc Guérin, Lara Urban

## Abstract

Wetland ecosystems provide interfaces for the transmission of microbial pathogens and antimicrobial resistances (AMR) between migratory birds, wild and domestic animals, and human populations. The efficient surveillance of wetlands is, however, challenging, since the typically low concentration of pathogens typically requires the sampling of large volumes of water and subsequent targeted detection, which is inherently limited to a few pathogens or AMR genes of interest. Here, we present a holistic, accessible, and cost-efficient framework to characterize the pathogen and resistance load of water sources together with their potential associated hosts by combining passive water sampling through torpedo-shaped devices with nanopore sequencing technology. We used this framework to characterize anthropogenically influenced and natural wetland ecosystems along the East Atlantic Flyway, where we obtained robust assessments of the microbial communities from long-read metagenomic and RNA virome data, and showed that anthropogenically impacted wetland ecosystems consistently exhibited higher relative abundances of pathogens and AMR genes. By focusing on avian influenza viruses (AIV), we finally highlight the additional need for targeted screening and whole-genome sequencing of pathogens of interest; we detected and characterized AIV at a third of the monitored sites, and used environmental DNA (eDNA) to explore potential animal hosts to better understand the role of wetland ecosystems as One Health interfaces.

**Graphical Abstract:** 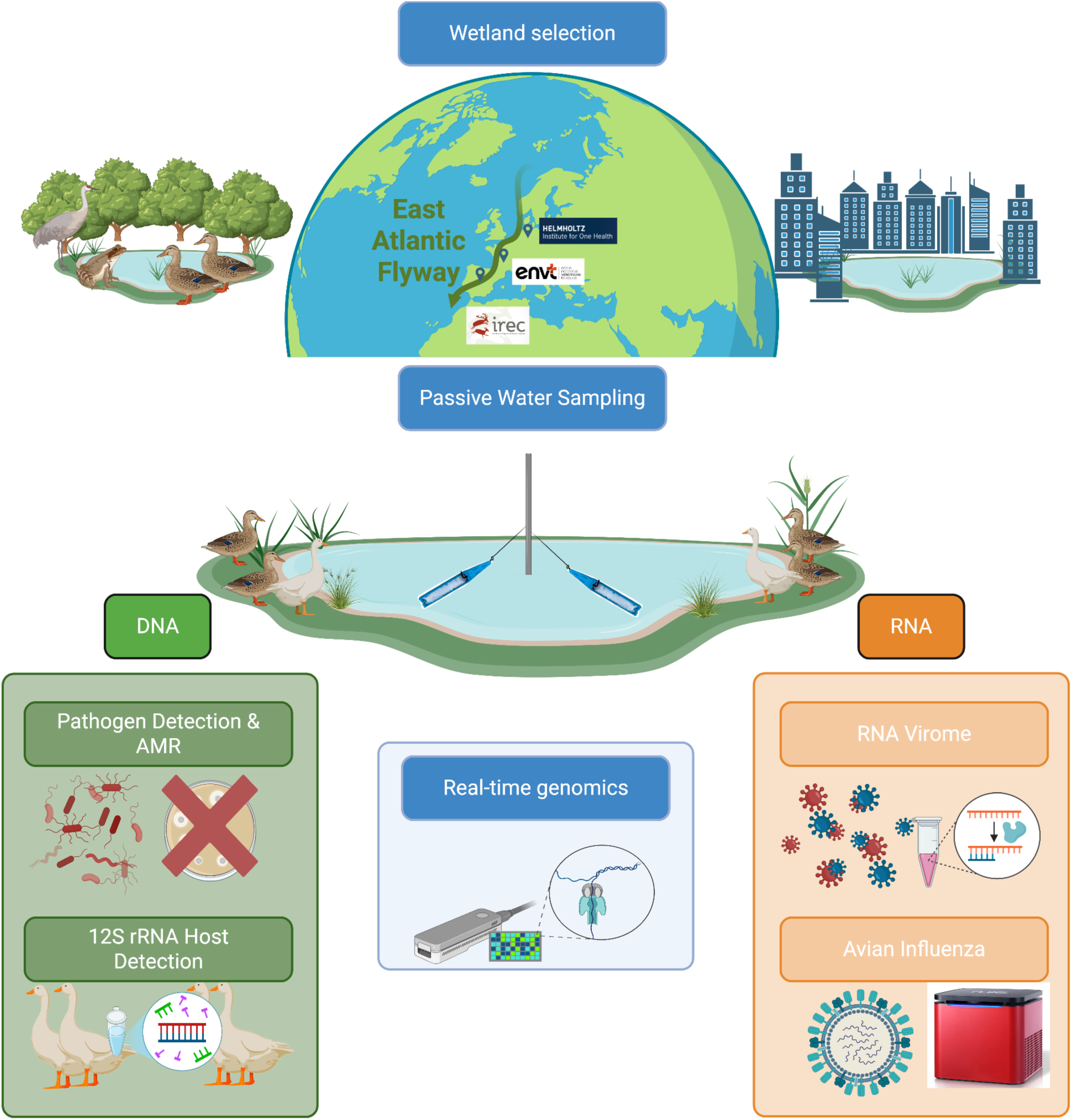

## 1. Introduction

Wetlands are among the most productive ecosystems on Earth, covering about 7% of the planet’s surface while supporting an estimated 40% of global biodiversity ^1^. They provide critical ecosystem services such as climate regulation, flood control, carbon sequestration, and water purification, and they serve as essential habitats for migratory birds ^2^. Despite this importance, wetlands are under severe threat from anthropogenic pressures, with Europe alone having lost more than half of its wetland area since 1970^3^. Their degradation not only undermines biodiversity conservation but also poses significant risks for public health, exemplifying the interconnectedness of environmental, animal, and human health as captured by the One Health concept ^4^.

Pollutants introduced by agriculture and urbanization, such as nutrients, pesticides, heavy metals, and antimicrobials, reshape wetland microbial ecology and foster environmental reservoirs of pathogens and antimicrobial resistances (AMR) ^5^. AMR, now recognized by the World Health Organization as one of the top ten global health threats ^6^, is driven in aquatic systems by the confluence of high microbial densities and sub-lethal antimicrobial concentrations, which facilitate horizontal gene transfer via mobile genetic elements ^7^. Resistance genes that emerge in such environments can spread to clinically relevant pathogens and subsequently re-enter animal and human populations, underscoring the environment’s role as a critical AMR reservoir ^5^.

In parallel, wetlands are hotspots for zoonotic pathogen transmission. RNA viruses are of particular concerns since they possess high evolutionary plasticity, enabling rapid host adaptation and establishing new transmission routes ^8^. RNA viruses also profit from global environmental changes, such as caused by increased livestock density, global mobility, and deforestation ^9^, which further increases the risk of the emergence of new zoonotic pathogens. Various RNA viruses such as *Picornaviridae*, *Paramyxoviridae*, or *Orthomyxoviridae* (such as Avian Influenza Viruses, or AIV) are known to impact the health of waterfowl in wetlands, and are therefore at risk of being globally distributed through migratory flyways ^10^. In the case of AIV, these avian hosts often asymptomatically carry low pathogenicity AIV (LPAIV), leading to spillover events to poultry and subsequent evolution into high pathogenicity AIV (HPAIV) with devastating animal welfare, economic, and food security consequences ^11^. Since the emergence of the H5N1 “Gs/GD lineage” in 1996, it has additionally become clear that HPAIV can also be directly transmitted from wild birds to poultry ^12^. Along the East Atlantic Flyway, recurrent outbreaks of H5N1 HPAIV have contributed to the ongoing panzootic in Europe ^13^, with severe consequences not only for poultry production but also for wildlife conservation, as increasing numbers of wild bird species have proven susceptible to the current lineages ^14,15^.

Traditional pathogen and AMR monitoring approaches from water ecosystems often rely on active filtration of large volumes of water, as in the case of AIV surveillance where the pathogen is expected to occur at low concentrations ^16^. Processing such large volumes is logistically challenging: field collection, preservation, and transport to the laboratory are difficult, and on-site filtration requires access to costly autosamplers and pumping equipment ^17^. Large-scale environmental surveillance, however, needs to be based on approaches that are inexpensive, simple to deploy and transport, and capable of integrating temporal and spatial signals. Recently developed 3D-printed, torpedo-shaped passive samplers meet these requirements: The torpedo design helps the sampler stay clean and prevents waterborne debris buildup, allowing its deployment for extended timeframes ^18^. The samplers can be easily assembled with a standard 3D printer, and deployed *in situ* at the point of care without specialized equipment or highly trained personnel. The torpedo-shaped passive sampler accommodates adsorptive media such as electronegative membranes to capture biological material during multi-day deployments ^18^. Originally optimized for wastewater surveillance of SARS-CoV-2, these devices have proven to be cost-effective, easy to install at scale, and sensitive to low pathogen concentrations, offering a practical alternative to grab sampling or autosampler-based approaches ^18^. Their simplicity and low cost have driven growing interest, and recent applications in urban wastewater have demonstrated effective surveillance not only of SARS-CoV-2, but also of fecal indicator bacteria and other respiratory viruses, including influenza A and B ^18–22^, as well as more recently AIV in wetland areas ^23^.

These studies have either relied on targeted approaches such as quantitative PCR detection or probe-based sequencing, which capture only a fraction of the microbial community, or on culture-based isolation ^24^, which can recover only a small proportion of the microbial diversity ^25^. To address these limitations, we here combined torpedo-shaped passive water samplers with long-read genomics by nanopore sequencing. Nanopore sequencing technology allows for holistic pathogen and AMR monitoring by integrating DNA-based metagenomics with RNA virome characterizations ^26–28^. The long nanopore reads allow for the analysis of extended contiguous sequences, which makes taxonomic and functional inferences more robust ^29^, elucidates the genomic context of resistance genes ^30,31^, and enable *de novo* assemblies for high-confidence pathogen and resistance gene detection ^32–34^. Whole RNA virome studies have revolutionized virus discovery, and have advanced our understanding of natural viromes in both vertebrates and invertebrates; recently, a nanopore sequencing-based protocol further facilitates the assembly of viral taxa by integrating long-read technology ^35–37^. Due to its capability to also sequence short reads as generated in metabarcoding studies, nanopore sequencing technology can further be leveraged for vertebrate environmental DNA (eDNA) analysis ^38,39^, to then identify potential wildlife or livestock hosts of the detected pathogens and resistances. We here apply this framework to characterize the pathogen load, the presence of AMR, and the potential vertebrate host range from passive water sampling of wetland ecosystems along the East Atlantic Flyway, and to assess the potential implications of anthropogenic pressures on wetlands for public health.

## 2. Methodology

### 2.1. Wetland selection

We sampled twelve lentic wetland sites across Germany, France, and Spain, selected to represent anthropogenic and natural land-use categories (Supplementary Table 1). Sites were coded using a [Country][Environment][Site Number] format, where Country is G (Germany), F (France), or S (Spain); and Environment is A (anthropogenic) or N (natural). The six anthropogenically impacted wetland sites were located adjacent to a landfill (GA1), to cities (GA2-3), to intensive duck farms (FA1-2), and to a sewage treatment plant (SA1). These were compared with six proximal natural sites, including coastal marshes and peat bogs (GN1-3) and protected wetland reserves (FN1; SN1-2). All sites were characterized by standing or slow-moving water (lentic systems), minimizing flow-related variability in sampler performance and ensuring comparable sampling conditions across locations. Detailed site descriptions, GPS coordinates, and environmental parameters are provided in Supplementary Table 1 and Supplementary Figure 1.

### 2.2. Passive water sampling

Passive water sampling was conducted using 3D-printed torpedo-shaped passive sampler units developed by the McCarthy group (Queensland University of Technology, Australia) ^18^. These torpedoes are designed to reduce clogging and fouling during deployment and can accommodate various passive sampling materials. For this study, each unit contained two 0.45 µm Mixed Cellulose Ester electronegative membranes (Whatman, Merck) for microbial particle adsorption. Two or three passive samplers were deployed at each wetland site. Each sampler was attached to a metal rod driven into the sediment using cable ties, positioning them 20 cm below the water surface to ensure continuous submersion while avoiding sediment contact. After three days in the water column, samplers were retrieved, visually inspected to remove debris, dismantled, and the membranes were transferred to bead-beating tubes for subsequent nucleic acid extraction.

### 2.3. Dual DNA and RNA extraction

From the bead-beating tubes with the electronegative membranes, we performed simultaneous extraction of DNA and RNA using the AllPrep PowerFecal Pro DNA/RNA kit (QIAGEN, 2018, Hilden, Germany), following the manufacturer’s protocol with modified elution volumes (25 µL DNA, 50 µL RNA). DNA and RNA yields were quantified using Qubit4.0 High Sensitivity assays (Invitrogen, 2021). Negative controls included field blanks, extraction blanks, and sequencing blanks, which were processed identically to the environmental samples.

### 2.4. Nanopore sequencing

Four parallel real-time genomic sequencing by nanopore sequencing approaches were employed: From DNA, we performed (i) shotgun metagenomics to characterise microbial communities and detect antimicrobial resistance genes; and (ii) 12S rRNA metabarcoding for vertebrate species detection; from RNA, we performed (iii) RNA virome analysis, and (iv) targeted detection of AIV by qPCR followed by whole-genome viral sequencing. Nanopore sequencing was performed on portable Oxford Nanopore Technologies (ONT, Oxford, UK) MinION Mk1c or Mk1d sequencers using R10.4.1 flow cells. Sequencing runs were conducted for 24 hours using the MinKNOW v24.11.10 software, with data acquisition at 5 kHz, a minimum quality score of 9, and a minimum read length threshold of 20 bases.

#### 2.4.1 Shotgun Metagenomics

Nanopore sequencing libraries from 24 samples (12 sites × 2 replicates) were generated using the Rapid Barcoding Kit (SQK-RBK114.24). To balance sequencing yield across samples, each sample was assigned two unique barcodes per sequencing run, with data from both barcodes combined during downstream analysis. To achieve equimolar pooling, samples were divided into two batches based on DNA concentration: eight high-concentration samples were pooled and distributed across two flow cells, and 16 lower-concentration samples plus sequencing blanks were pooled separately and distributed across another two flow cells, following established protocols ^33^. One negative sequencing control was included per flow cell. One sampling and two extraction negative controls were sequenced separately.

#### 2.4.2 Vertebrate Metabarcoding

We PCR-amplified a ∼97 base pair (bp) fragment of the mitochondrial 12S rRNA gene (V5 region) from the eDNA extracts from our 12 locations, one technical negative control, one negative control of extraction, and one positive control following our previously established protocol ^39^. Briefly, We used the broad-vertebrate primer set 12SV05 (forward 5′-TTAGATACCCCACTATGC-3′, reverse 5′-TAGAACAGGCTCCTCTAG-3′) developed by Riaz *et al.* (2011) ^40^. Unique 9 bp tags (with ≥3 bp differences between any two tags) were appended to both primers for sample multiplexing. PCR mix (20 µL volume) contained 3 µL of template DNA, 0.75 U AmpliTaq Gold polymerase, 1× Gold Buffer, 2.5 mM MgCl₂, 0.2 mM each dNTP, 0.6 µM of each tagged primer, and 0.5 mg/mL bovine serum albumin. To suppress human DNA amplification, a 3 µM human-blocking oligonucleotide (5′-TACCCCACTATGCTTAGCCCTAAACCTCAACAGTTAAATC-3′, with a 3′ C3 spacer) was included in each reaction, following ^41^. The PCR thermal profile was initial denaturation at 95 °C for 10 min; 45 cycles of 94 °C for 30 s, 60 °C for 30 s, 70 °C for 60 s; and a final extension at 72 °C for 7 min. Successful amplification was confirmed on 2% agarose E-Gel™ gels (ThermoFisher). Amplicon pools from each sample were then combined (18 µL per PCR), purified with HigherPurity™ magnetic beads (1:1 ratio) and eluted in 50 µL of DNA-free water. One sequencing library was prepared from the purified 12S rRNA amplicon pool using the Ligation Sequencing Kit (SQK-LSK114) in one flow cell.

#### 2.4.3 RNA Virome

RNA extracts were processed following the Rapid SMART-9N protocol described by^27^, which uses random priming and a template-switching mechanism to generate long-length cDNA flanked by known sequences for downstream PCR barcoding. Briefly, residual DNA was removed from 44 µL RNA by treatment with 5 µL TURBO DNase buffer and 1 µL TURBO DNase (Thermo Fisher Scientific) at 37 °C for 30 min. RNA was then purified and concentrated to 10 µL using the RNA Clean & Concentrator-5 kit (Zymo Research). For first-strand cDNA synthesis, 10 µL RNA was mixed with 1 µL RLB RT 9N primer (5′-TTTTTCGTGCGCCGCTTCAACNNNNNNNNN-3′, 2 µM), a random-priming oligonucleotide with a fixed 5′ sequence complementary to the PCR primers, and 1 µL of 10 mM dNTPs, incubated at 65 °C for 5 min, and placed on ice. Eight microlitres of this mixture were reverse transcribed in a 20 µL reaction containing the annealed RNA, 4 µL SuperScript IV First-Strand Buffer, 1 µL 0.1 M DTT, 1 µL RNaseOUT, 1 µL RLB TSO oligo (5′-GCTAATCATTGCTTTTTCGTGCGCCGCTTCAACATrGrGrG-3′, 2 µM), which hybridises to non-templated cytosines added by the reverse transcriptase and adds a second fixed sequence at the cDNA 5′ end, and 1 µL SuperScript IV reverse transcriptase (Thermo Fisher Scientific). The reaction was incubated at 42 °C for 90 min followed by 70 °C for 10 min. The resulting double-tagged cDNA was amplified in a 50 µL PCR reaction containing 5 µL cDNA, 25 µL LongAmp Taq 2x Master Mix (New England Biolabs), 19.5 µL nuclease-free water, and 0.5 µL RLB barcode primer (SQK-RPB114.24), which binds the fixed primer sites at both cDNA ends and incorporates a sample-specific barcode. PCR cycling conditions were: 95 °C for 45 s; 25–30 cycles of 95 °C for 15 s, 56 °C for 15 s, and 65 °C for 5 min; final extension at 65 °C for 10 min. Amplicons were purified using a 1:1 AMPure XP bead ratio (Beckman Coulter) and quantified with a Qubit 4.0 fluorometer using the dsDNA High Sensitivity Assay Kit (Thermo Fisher Scientific). Sequencing libraries were prepared with the Rapid Sequencing PCR Barcoding Kit (SQK-RPB114.24) and sequenced on two flow cells.

#### 2.4.4. AIV analysis

AIV detection was performed by applying reverse transcription(RT)-qPCR to the extracted RNA while targeting a highly conserved region of the influenza matrix 1 (M1) gene with a one-step Taqman RT–PCR in a Fast7500 equipment (Applied Biosystems), using primers (M+ 25 5’-AGATGAGTCTTCTAACCGAGGTCG-3’, M-124 5’-TGCAAAAACATCTTCAAGTCTCTG-3) and probe (M+ 64 FAM 5’ TCAGGCCCCCTCAAAGCCGA-3’TAMRA) ^42^ and adding bovine serum albumin to increase sensitivity and avoid PCR inhibitors ^43^. Whole-genome AIV sequencing of positive samples was performed after complementary DNA (cDNA) conversion and multi-segment amplification using M-RTPCR ^44^, targeting conserved regions across all AIV segments: Briefly, the RNA was mixed with SuperScript III One-Step PCR (Invitrogen) buffer and enzyme mix (containing the reverse transcriptase enzyme and the PCR amplification enzyme). The thermal cycle parameters were 42 °C for 60 min, 94 °C for 2 min, and then 5 cycles (94 °C for 30 s, 45 °C for 30 s, and 68 °C for 3 min), followed by 31 cycles (94 °C for 30 s, 57 °C for 30 s, and 68 °C for 3 min). Primers used were MBTuni-12 (5′-ACGCGTGATCAGCAAAAGCAGG) and MBTuni-13 (5′-ACGCGTGATCAGTAGAAACAAGG) that correspond to the 5′ and 3′ conserved sequences of all eight influenza A segments, and that have been demonstrated effective on all subtypes. We finally used the Rapid Barcoding Kit (SQK-RBK114.24) ^44^ and sequenced the library on one flow cell.

### 2.5. Bioinformatic analysis

#### 2.5.1. Nanopore data processing

Raw nanopore signal data (POD5) from all sequencing runs were basecalled using Dorado v5.0.0 in super-accuracy mode (dna_r10.4.1_e8.2_400bps_sup@v5.0.0; ^45^). Demultiplexing and adapter/barcode trimming for shotgun and virome reads were performed with Porechop (v0.2.4) ^46^. Reads shorter than 100 bases (150 bases for AIV sequencing analysis) and reads with quality score below 9 (quality score below 8 for AIV sequencing analysis) were discarded using NanoFilt (v2.8.0) ^47^.

#### 2.5.2. Metagenomic taxonomic classification

Quality-filtered nanopore reads were taxonomically classified using Kraken2 v2.1.2 ^48^ against the NCBI nt_core database (May 2025). For community composition analyses, all water samples were downsampled to 87,000 reads using SeqKit v2.3.0 ^49^. Sample GA3.1 (28,000 reads) was excluded due to insufficient sequencing depth. Beta-diversity between samples was quantified using Bray-Curtis dissimilarity on genus-level reads and visualized by principal coordinate analysis (PCoA). Up to 20 most abundant microbial genera at a minimum relative abundance of 1% were visualized with stacked bar plots generated in Python (v3.12) using the pandas and matplotlib libraries, with other taxa grouped into the “Others” category.

*De novo* assembly was performed using metaFlye (v2.9.6) ^50^ and nanoMDBG (v1.1) ^51^ to identify the optimal approach for our dataset. The metaFlye assemblies underwent a polishing workflow: reads were first realigned to the assemblies using minimap2 (v2.28) ^52^, followed by three iterative rounds of polishing with Racon (v1.5.0) ^53^ and Medaka (v1.7.2) ^54^. NanoMDBG assemblies were polished with Medaka (v1.7.2) alone. Assembled contigs were subsequently annotated using Kraken2 for taxonomic classification and Prokka v1.14.5 for functional annotation ^55^.

#### 2.5.3. Metagenomic pathogen and virulence analyses

For stringent pathogen identification, reads and contigs were aligned to the NCBI nt_core databases using minimap2 (v2.28) and processed with MEGAN-CE v6.21.1 ^56^, using the lowest common ancestor (LCA) assignment. Taxonomic assignment was only accepted if more than 50% of near-best alignments for a read matched the same genus. This conservative approach, combined with a ensures high-confidence species-level assignments by classifying ambiguously mapped sequences at higher taxonomic ranks. Pathogenic species were subsequently identified using the Chan Zuckerberg ID initiative pathogen species list ^57^. Only pathogenic species with at least 5 reads in a given replicate (or sampling site) were considered for downstream analysis.

To further investigate the pathogenic potential of *Vibrio cholerae*, targeted screening for specific virulence factors was conducted: Alignments against the core protein dataset of the Virulence Factor Database (VFDB) ^58^ were performed using DIAMOND BLASTx (v2.1.13) ^59^ to specifically search for reads corresponding to the cholera toxin subunit genes, *ctxA* and *ctxB* ^58^.

AMR and virulence genes were detected using AMRFinderPlus v4.0.23 ^60^ in --plus mode on the complete dataset with its default thresholds (including minimum sequence identity of 90%). For assembled contigs, open reading frames were predicted with Prodigal v2.6.3 ^61^, enabling nucleotide- and protein-level analyses with AMRFinderPlus to maximize sensitivity. Pathogen-AMR associations were considered high-confidence when the same resistance gene was detected on both reads and contigs from a sample, and when the same reads/contigs were taxonomically linked to the same pathogen.

To assess the mobility potential of detected AMR genes, assembled contigs were screened for plasmids using PlasmidFinder v2.1.6 ^62^ against the PlasmidFinder database.

#### 2.5.4. Avian host detection from eDNA

Basecalled 12S rRNA amplicon reads were demultiplexed by their 9 bp primer tags using OBITools4 obimultiplex (version 1.3.1) ^63^, allowing up to two mismatches in tag recognition. Primer sequences were trimmed from reads with Cutadapt (v4.2) ^64^. The reads were then processed in VSEARCH (v2.21) ^65^: we filtered reads by expected error (maxEE 1.0) to remove low-quality reads, dereplicated reads (discarding singletons), removed chimeras, and clustered the remaining sequences into operational taxonomic units (OTUs) at 97% similarity. Each OTU representative sequence was taxonomically identified by global alignment against the MIDORI2 reference database of mitochondrial sequences (version “Unique 266”) ^66^. Assignments were limited to bird species (Aves class), and accepted as robust if an alignment covered ≥80% of the query, had ≥98% identity, and if a taxonomic assignment had at least five reads as hits after OTU aggregation of same species detection. When species-level calls were biogeographically implausible for our sampling sites (i.e., the species is not present in the sampled area), we collapsed the assignment to a higher rank (i.e., genus or family) or removed the record if no meaningful higher-rank assignment was possible. Regional occurrence (presence/absence) was assessed with eBird (https://ebird.org/home). Multiple OTUs mapping to the same retained taxon within a sample were summed to produce per-taxon read counts.

#### 2.5.5. RNA Virome taxonomic analysis

*De novo* assembly was performed using nanoMDBG (v1.1) and polished with Medaka (v1.7.2). Assembled contigs were compared against the NCBI non-redundant protein database (NR database, accessed May 2025) using DIAMOND BLASTx (v2.1.13) ^59^. Contigs with a percentage identity above 80% when matched to the kingdom “Viruses” (NCBI taxid: 10 239) were annotated as viral contigs.

#### 2.5.6. AIV whole-genome sequencing analysis

The whole-genome AIV sequencing FASTQ files were aligned to the reference genomes using minimap2 (v2.28) ^52^ with the -ax map-ont setting. We used a reference database generated for each segment from the NCBI Influenza Virus Database, which contains all AIV nucleotide sequences from Europe (as of 04/03/2023). The resulting SAM files were converted to BAM format, sorted, and indexed using SAMtools (v1.17) ^67^. Using samtools idxstats, we selected the segment reference to which most reads mapped across every segment. All our reads were then mapped to the best reference for each of the eight segments of the influenza genome using minimap2 (v2.28) and genome coverage distributions were obtained. We then obtained the final AIV consensus sequence with BCFtools (v1.17) ^68^. For subtyping of H4 hemagglutinin (HA) consensus segment we used GISAID BLAST tool ^69^ and FluSurver (http://flusurver.bii.a-star.edu.sg).

To contextualize the H4 HA consensus sequence, we downloaded all publicly available full-length H4 HA sequences from GISAID since 2015 (accessed the 25^th^ of August 2025). Metadata (host, location, collection date) were curated from the associated GISAID isolate tables. To reduce redundancy and improve tree readability, we implemented a stratified subsampling approach: sequences were grouped by host, country, and year, and up to three representatives per group were retained randomly. Multiple sequence alignment was performed with MAFFT (v7.526) ^70^. The aligned dataset was used to infer a maximum-likelihood phylogeny with IQ-TREE2 (v2.3.4) ^71^ under the best-fit nucleotide substitution model selected by ModelFinder Plus (-m MFP). Node support was assessed with 1,000 ultrafast bootstrap replicates (-bb 1000) and 1,000 SH-aLRT replicates (-alrt 1000). The resulting tree was visualized in iTOL (v7) ^72^.

## 3. Results

### 3.1. Nanopore sequencing data

After dual-nucleotide extraction of the electronegative membranes from the torpedo-shaped passive samplers (Methodology), DNA yields ranged from below detection limit to 918 ng (Supplementary Table 2), while RNA yields ranged from below detection to 3,335 ng (Supplementary Table 3; Supplementary Figure 2A). Metagenomic sequencing generated 28k to 863k reads after filtering (median lengths 485-1,534 bp). 12S rRNA amplicon sequencing generated 19M reads after filtering. RNA virome sequencing produced 87k to 2.1 M reads (median lengths 375–1,663 bp; Supplementary Table 3; Supplementary Figure 2B-E). AIV sequencing generated 600k reads after filtering (median length 323).

### 3.2. Microbial taxonomic compositions across wetland sites

Taxonomic classification success varied considerably, with 13k–365k (corresponding to 16-54%) metagenomic reads assignable to microbial genus level. After downsampling to 87k reads for comparative analysis, one sample (GA3.1) was excluded due to insufficient reads (Supplementary Figure 2F). All negative controls resulted in low DNA yields (<0.1 ng). Sampling negative controls included read mapping to the *Homo*, *Sphingomonas*, and *Timema* genera. Extraction negative controls included reads mapping to *Homo*, *Mus*, and *Timema* genera. Sequencing negative controls included reads mapping to the *Homo*, *Timema*, *Photobacterium*, and *Rheinheimera* genera. All pathogen-containing genera were absent from all negative controls (Supplementary Table 2).

The microbial communities on the taxonomic genus level per site were highly similar in the respective biological replicates but otherwise did not show any clustering according to land usage or country of origin (Figure 1A). Within the German sites, anthropogenic samples exhibited higher relative abundances of *Paracoccus* and *Pseudomonas*, whereas natural sites showed increased *Marivivens* and *Curvibacter* (Figure 1B). French sites displayed distinct, site-specific profiles: FA1 was dominated by *Aeromonas*, FA2 by *Streptomyces*, and FN1 by *Hydrogenophaga* (Figure 1C). The Spanish anthropogenic site SA1 was enriched for *Vibrio* and *Aeromonas*, while natural sites showed higher *Streptomyces* and freshwater-associated taxa, including phototrophs such as *Cyanobium* and *Limnohabitans* (Figure 1D).

**Figure 1.**
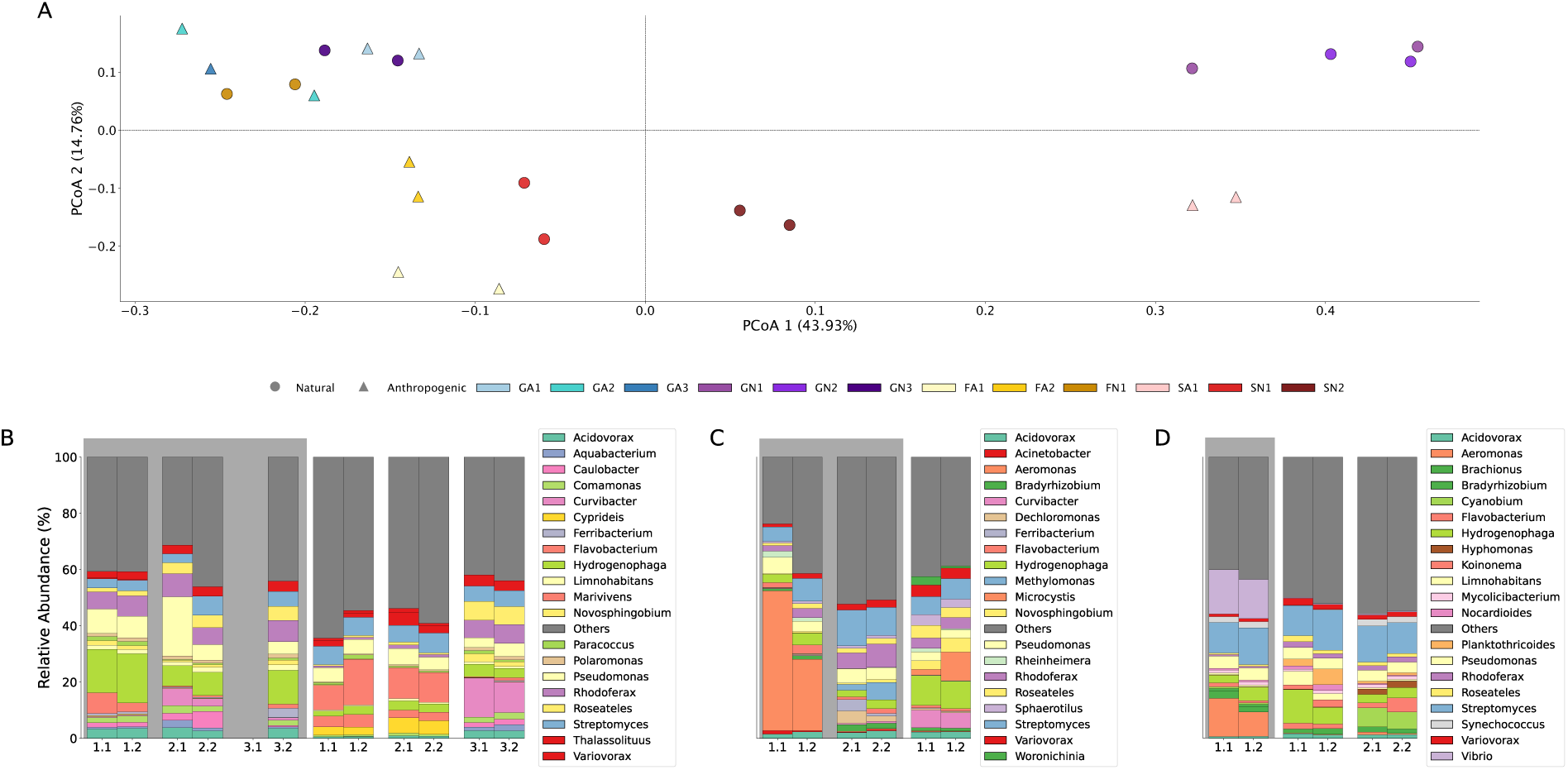
Microbial taxonomic composition on the genus level across wetland sites. (after downsampling; two biological replicates per sample; see Methodology). **(A)** Principal Coordinate Analysis (PCoA) of Bray-Curtis dissimilarities. Sites were coded using a [Country][Environment][Site Number] format, where Country is G (Germany; blue-purple gradient), F (France; yellow gradient), or S (Spain; red gradient); and Environment is A (anthropogenic; triangular shape) or N (natural; circular shape). **(B–D)** Relative abundance of the 20 most abundant bacterial genera for **(B)** Germany (including one replicate dropout after downsampling), **(C)** France, and **(D)** Spain. The grey background highlights anthropogenically impacted sites. Genera outside the top 20 are aggregated as “Others” (dark grey).

### 3.3. Microbial pathogen and antimicrobial resistance detection across wetland sites

Stringent microbial pathogen species assignment detected the presence of pathogens in all wetland sites (Supplementary Table 4; Methodology). The anthropogenically impacted sites contained 9,346 total pathogenic reads across 17 unique pathogen species (≥5 reads threshold), while natural sites contained 703 pathogenic reads across 15 unique species. The French duck farm (FA1) showed the highest pathogen diversity and relative abundance, with the presence of several pathogenic *Aeromonas* species: *A. veronii* (2,960 and 656 reads in FA1.1 and FA1.2), *A. hydrophila* (223 and 481 reads), *A. caviae* (81 and 45 reads) and also *Pseudomonas putida* (212 and 189 reads) (Figure 2). The Spanish anthropogenic site (SA1) had 1,809 reads mapping to *Vibrio cholerae*, with co-detection of *A. veronii* (139 reads) (Figure 2).

**Figure 2.**
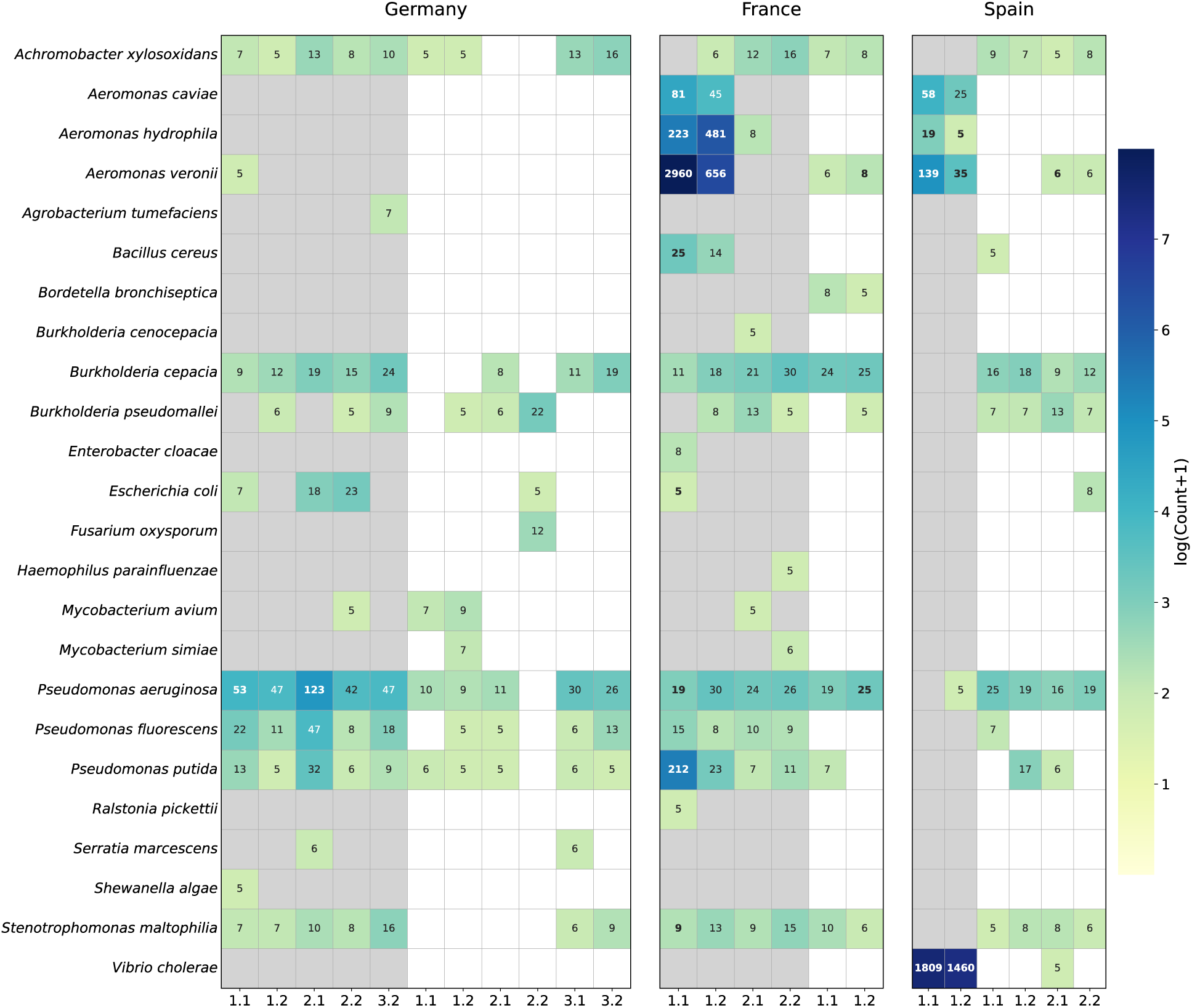
Number of sequencing reads assigned to pathogenic bacterial species across wetland sites. (after downsampling; two biological replicates per sample; see Methodology). The grey background highlights anthropogenically impacted sites. The reads assigned to pathogenic species that were additionally identified by *de novo* assemblies are highlighted in bold.

Resistome analysis on the sequencing read level led to the detection of several AMR genes in both biological replicates taken from the anthropogenically impacted French (FA1) and Spanish (SA1) wetland sites (Table 1; Methodology). The French wetland FA1 that was adjacent to a duck farm contained nine distinct AMR genes, dominated by beta-lactamases conferring resistance to β-lactam antibiotics: *bla*_OXA_ (extended-spectrum β-lactamase), *cphA* and *cphA1* (metallo-β-lactamases conferring carbapenem resistance), and *bla*_FOX_ (AmpC-type cephalosporinase). The Spanish wetland SA1 harbored twelve unique AMR genes, including *bla*_VCC-1_, an Ambler class A carbapenemase. Sulfonamide resistance genes (*sul1*, *sul2*), which confer resistance to sulfonamide antibiotics commonly used in agriculture, were detected at both FA1 and SA1. While the GN3.2 sample contained one AMR gene, the *bla*_CAU_ gene, we excluded this sample from subsequent AMR analyses since this gene has been described as an intrinsic gene in environmental *Caulobacter crescentus* ^73^.

**Table 1.**
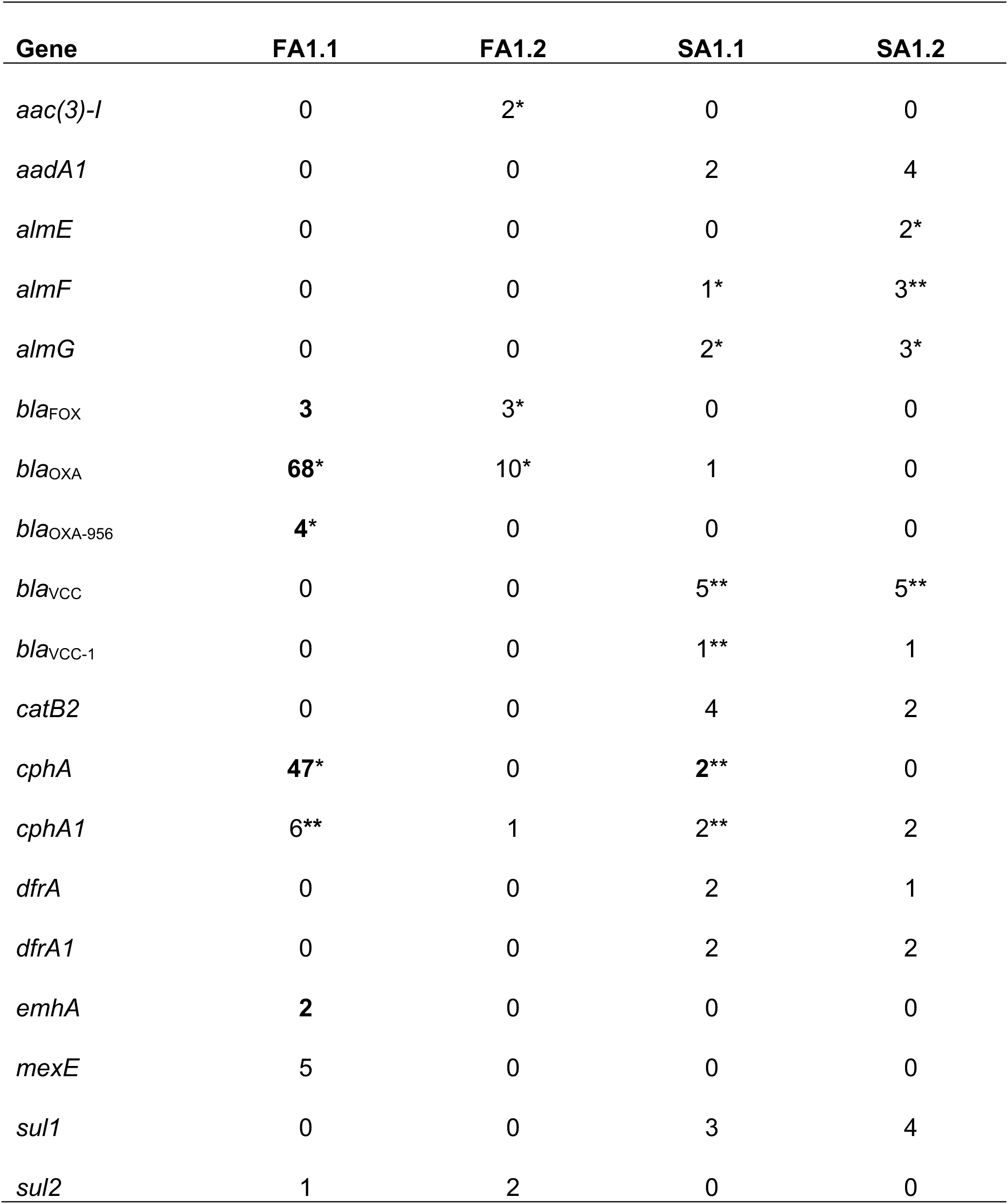
Number of detected antimicrobial resistance (AMR) genes across AMR-positive wetland sites. AMR detection was done on the sequencing read-level using AMRFinderPlus; all genes that had at maximum one hit across all samples were excluded. A single asterisk (*) indicates AMR gene counts that could be assigned to pathogenic bacterial species; a double asterisk (**) indicates AMR gene counts that could be assigned to a bacterial genus that is likely pathogenic; bold AMR gene counts were confirmed by detection on *de novo* assemblies of pathogenic bacterial species in the same sample (Methodology). The GN3.2 sample is excluded since it only contains *bla*_CAU_ as an intrinsic gene in environmental *Caulobacter crescentus*.

We used the taxonomic assignment of AMR gene-containing sequencing reads to link AMR genes to pathogenic microbial species (Table 1). At FA1, we found that the *bla*_OXA-956_, *bla*_OXA_, and *cphA* genes were encoded by the pathogen *Aeromonas veronii*. To further assess the pathogenic potential of taxa carrying AMR genes beyond our stringent pathogenic species annotation criteria, we expanded our analysis to the genus level in the cases where the detected genus was predominantly pathogenic in the same sample (Methodology). At SA1, *bla*_VCC_ and *bla*_VCC-1_-carrying reads were assigned to the *Vibrio* genus, and most Vibrio-assigned reads in the same sample could be classified as the pathogenic *Vibrio cholerae*. Similarly, *cphA*- and *cphA1*-carrying reads were assigned to the *Aeromonas* genus, and most reads in the same sample could be classified as the pathogenic *A. veronii*, *A. hydrophila*, and *A. caviae* species (Table 1).

To validate pathogen detections and pathogen-AMR associations at increased confidence, we repeated our analyses on the *de novo* assembly-level (see bold numbers in Figure 2; Table 1). While metaFlye generated assemblies with superior contiguity metrics, nanoMDBG yielded greater total assembly size, contig count, taxonomic diversity, and gene content (Supplementary Figure 2; Supplementary Table 2; Methodology). Based on nanoMDBG assemblies, anthropogenic and natural sites yielded 892 and 5 contigs assigned to pathogenic species, respectively. At FA1, 191 contigs were assigned to *Aeromonas veronii* (Supplementary Table 5), with multiple contigs carrying β-lactamase genes previously identified in the read analysis: three contigs harbored *cphA* metallo-β-lactamase genes, three carried *bla*_OXA_ genes, and one contained the *bla*_OXA-956_ variant (Table 2). Additionally, two contigs that were assigned to *Escherichia coli* contained the *bla*_FOX_ β-lactamase (Table 2). At SA1, 235 contigs mapped to *Vibrio cholerae*, and these contigs lacked the typical cholera toxin genes *ctxA* and *ctxB* (Supplementary Table 5; Methodology). The *de novo* assemblies additionally revealed previously undetected AMR genes: At FA1, contigs classified as *Acidiphilium multivorum* harbored arsenic resistance genes *arsB* and *arsC*, and one contig classified as *Acinetobacter pittii* harbored the nickel-resistance gene *nreB* (Table 2). No plasmids were detected in any sample.

**Table 2.**
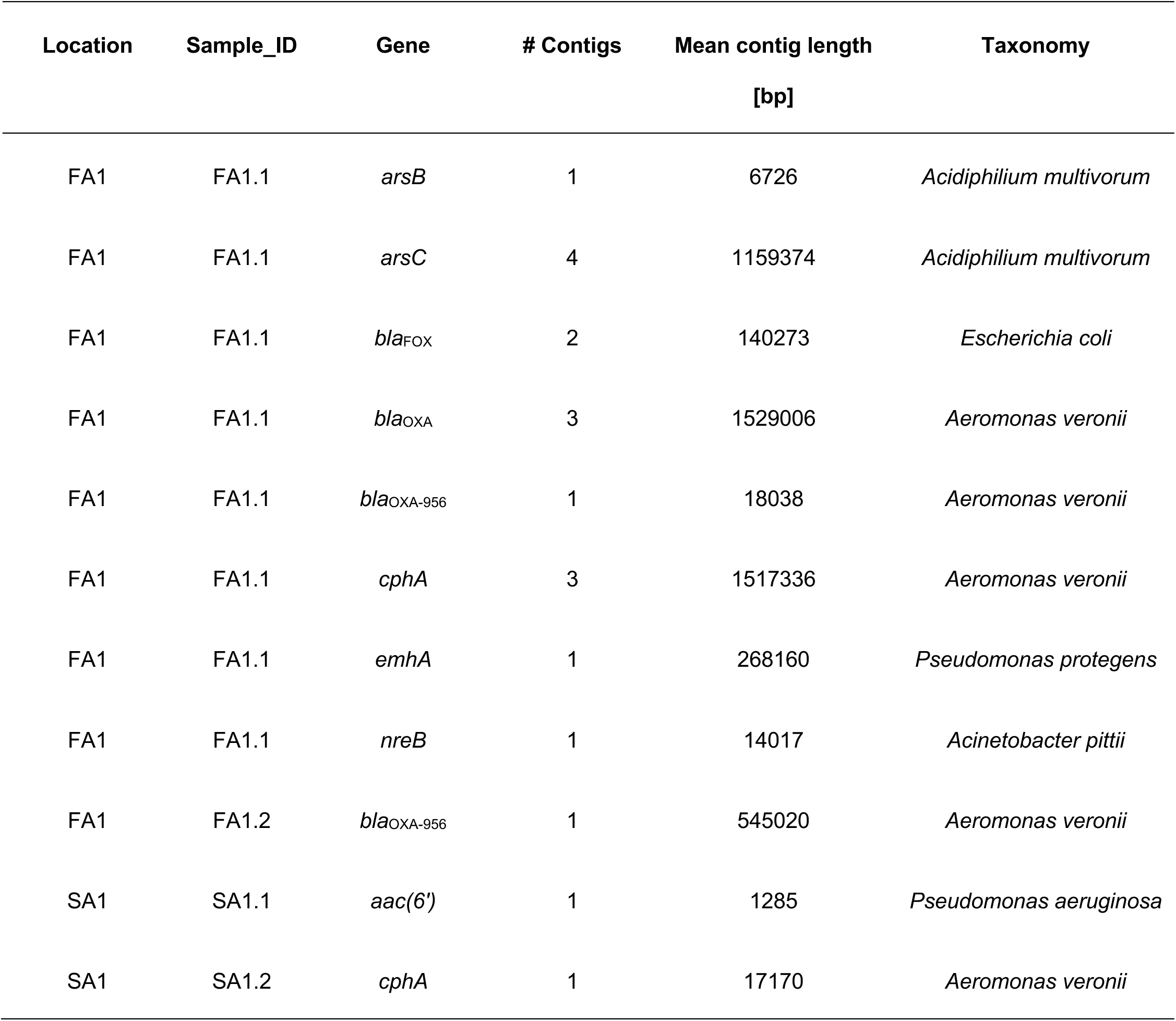
Resistance gene detection on the assembly level. For each antimicrobial and resistance gene, the wetland site, the biological replicate, the number of assembled contigs, the mean length of those contigs in base pairs (bp), and the taxonomic assignment of those contigs are indicated.

### 3.4. eDNA-based avian host inference

We detected avian eDNA using 12S rRNA metabarcoding at most of the wetland sites, except for GN1-3 and GA2 (Supplementary Figure 3; Methodology). Overall, detections were dominated by common waterfowl birds, with dabbling ducks (*Anas*, *Spatula*, *Mareca*), geese (*Anser*), swans (*Cygnus*), and cormorants (*Phalacrocorax*) being the most common taxa (Figure 3). The species richness (species per location) was highest at SN1 and FA1, followed by FA2 and SA1 (Figure 3). No avian taxa were detected in any negative control.

**Figure 3.**
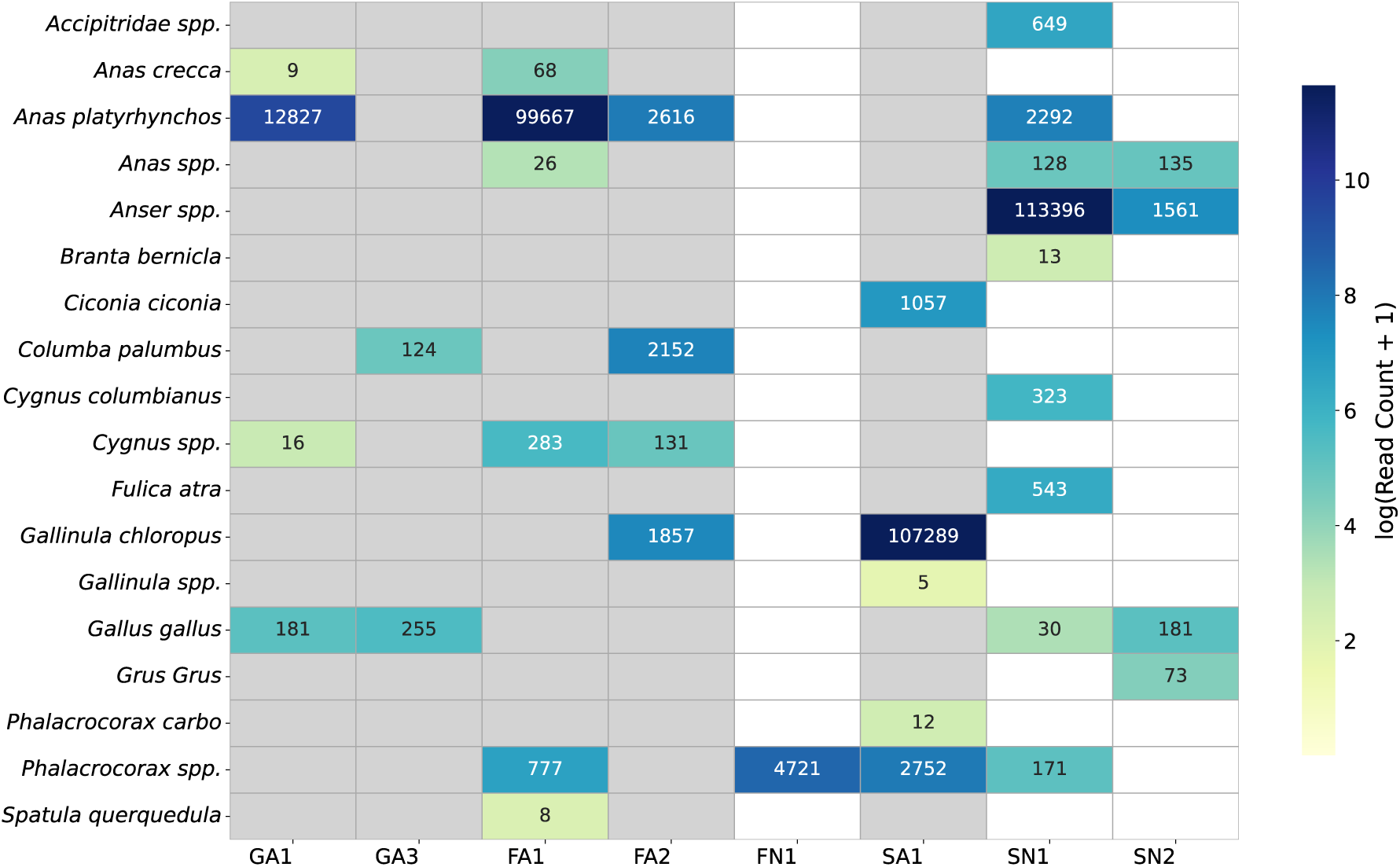
Number of avian species identified from 12S rRNA-based eDNA across wetland sites. Color intensity corresponds to log(read count + 1) aggregated per species and site. Species assignments were restricted to avian species at an identity percentage to the reference of ≥ 98% and read count of ≥ 5. The grey background highlights anthropogenically impacted sites.

### 3.5. RNA virome analysis

The percentage of sequencing reads assigned to the viral kingdom after random amplification of extracted RNA ranged from 0.1% to 1.26% (Supplementary Table 3; Methodology). After *de novo* assembly (Methodology), the contigs assigned to the viral kingdom ranged from none (FA1, FA2, and SA1) to 4.54% at GA3 (Supplementary Table 3). Due to unspecific taxon classification at the species and genus level, we focused our viral taxonomic classification on the family level. We overall found 93 contigs assigned to viral families, ranging from 0 contigs to 29 contigs per site (Supplementary Table 3), and covering 13 distinct viral families (Figure 4). The *Dicistroviridae* (20 contigs) and *Fiersviridae* (19 contigs) viral families were the most common ones (Figure 4). The *Picornaviridae* viral family, which is the only one that contains several potential mammalian and bird pathogens, was only detected by a few contigs (Figure 4). Viral contig recovery was highest at GA3 (29 contigs), followed by GN2 (17 contigs; Figure 4).

**Figure 4.**
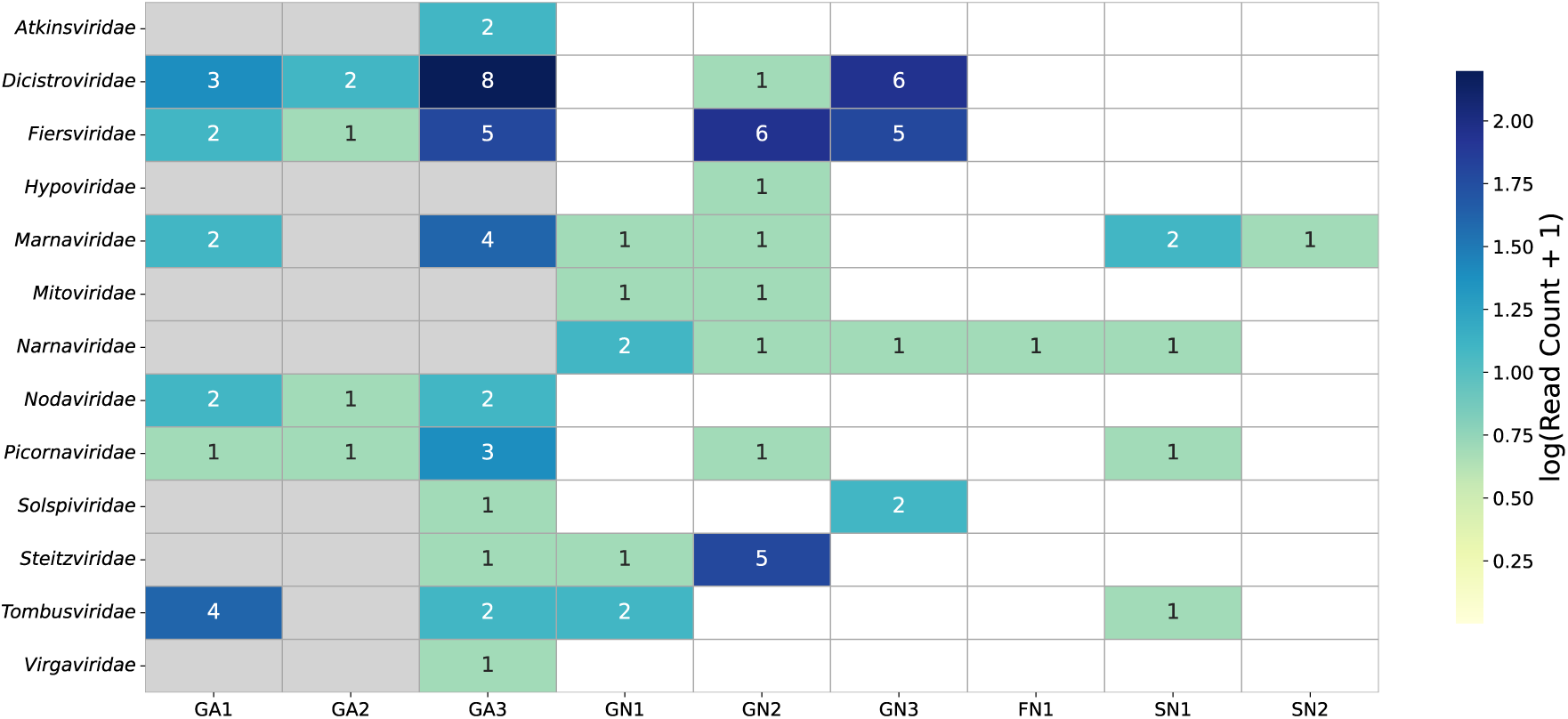
Number of viral family detections from *de novo* assemblies across wetland sites. Color intensity encodes log(contig count + 1) (Methodology). The grey background highlights anthropogenically impacted sites.

### 3.6. AIV detection and sequencing

We detected AIV from passive samplers at four of the 12 sampled wetland locations (Table 3), including all three wetland sites in France which were located in the vicinity of duck farms, and one wetland site in Spain, which was located close to a national park. The qPCR cycle threshold (Ct) values of the positive samples ranged from 34 to 40, with 40 representing the limit of detection (Table 3). HA subtyping by multi-segment PCR and whole-genome sequencing was successful only for one sample from site FA2. The HA consensus sequence had an average coverage depth of 160x and was classified as subtype H4, corresponding to a low pathogenicity AIV (LPAIV). We built a phylogenetic tree to place this strain in the evolutionary context of H4 viruses detected in Europe over the past ten years (Supplementary Figure 4).

**Table 3.** Detection and sequencing of avian influenza virus (AIV) across wetland sites. Positive detections were confirmed by RT-qPCR (Ct values shown). HA subtyping was performed where possible by multi-segment PCR and whole-genome sequencing. Potential avian hosts were inferred by combining results from vertebrate eDNA analyses by 12S rRNA metabarcoding with official reports of AIV in wild birds.

| Site | Positive / total<br>replicates | Ct | HA subtype | potential avian host |
| --- | --- | --- | --- | --- |
| FN1 | 1/6 | 37 | NA | *Phalacrocorax spp.* |
| FA1 | 4/5 | 34,36,37,40 | NA | *Anas platyrhynchos* (mallard), *Anas crecca* (teal), *Spatula querquedula* (garganey), *Cygnus spp.* (swans), *Phalacrocorax spp.* (cormorants), |
| FA2 | 2/3 | 37,38 | H4 | *Anas platyrhynchos* (mallard), *Gallinula chloropus* (moorhen), *Columba palumbus* (wood pigeon), *Cygnus spp.* (swans) |
| SN2 | 1/3 | 39 | NA | *Gallus gallus* (chicken), *Anser spp.* (geese), *Grus Grus* (cranes) |

To identify potential avian hosts associated with AIV-positive wetlands, we overlapped our vertebrate eDNA-based species detections (Figure 3) with official reports of AIV cases in wild birds ^74^. This revealed that AIV detections coincided with the detection of several potential host species (Table 3); for example, at wetland sites adjacent to duck farms (FA1, FA2), AIV detection coincided with the detection of multiple dabbling duck species (*Anas platyrhynchos, Anas crecca, Spatula querquedula*), which are known AIV reservoirs, as well as *Cygnus spp.* (swan) and *Phalacrocorax spp.* (cormorant) species (Table 3). At FA2, where AIV subtyping confirmed an H4 LPAIV, vertebrate eDNA also detected the presence of *Gallinula chloropus* (moorhen) and *Columba palumbus* (wood pigeon) (Table 3).

## 4. Discussion

Wetlands along the East Atlantic Flyway are One Health interfaces where migratory birds, domestic animals, and human populations converge, facilitating the circulation of pathogens and antimicrobial resistance across countries ^75,76^. Protecting and monitoring these ecosystems is therefore essential not only for biodiversity conservation but also for early warning of transboundary health threats at the wildlife– livestock–human interface. We here present a holistic, accessible, and cost-efficient framework to characterize the pathogen and resistance load of water sources together with their potential associated hosts by combining passive water sampling with nanopore sequencing. We then apply this framework to characterize anthropogenically influenced and natural wetland ecosystems along the East Atlantic Flyway, with *de novo* assemblies confirming their presence and enabling direct links between AMR genes and specific pathogens. RNA virome analysis revealed diverse viral families and highlighted both the potential and the current limitations of untargeted viral metagenomics—as highlighted by our subsequent targeted detection and sequencing of AIV, which was not identified by the RNA virome analyses. Finally, eDNA metabarcoding provided inference of potential AIV hosts, offering potentially valuable insights into the wildlife–livestock interface.

The microbial community could be robustly assessed across different wetland sites from metagenomic data obtained from the deployed torpedo-shaped passive sampling devices: The high similarity of the microbial composition at the taxonomic genus level between biological replicates demonstrates the site-specific replicability—similar to what we have previously observed from metagenomics from active air samples ^33^. Our comparison of microbial *de novo* assembly tools revealed that while the established metaFlye tool yielded more contiguous assemblies ^50^, the novel nanoMDBG assembler^51^ captured more biological insights, including more diverse microbial species annotations and improved gene recovery. This might be explained by metaFlye’s default filter discarding reads below 1kb, which comprise over 50% of our data; this highlights the advantage of nanoMDBG when it comes to fragmented environmental _DNA_ ^50,77^.

The metagenomic data contained several pathogens on the read- and assembly-level analysis, despite stringent pathogen filtering criteria ^78^. The anthropogenically impacted wetlands harbored over 13-fold more pathogen-associated reads (9,346 reads across 17 species) compared to natural wetland (703 reads across 15 species). *De novo* assemblies confirmed the presence of potentially clinically relevant bacteria such as *Acinetobacter baumannii*, *Pseudomonas aeruginosa*, and *Stenotrophomonas maltophilia*, and members of the Enterobacteriaceae family (*Citrobacter freundii*, *Escherichia coli*, *Klebsiella pneumoniae*). Additionally, we were able to generate assemblies of the pathogens *Bacillus cereus*, *Vibrio cholerae*, *Aeromonas caviae*, *Aeromonas hydrophila*, *Aeromonas veronii*, *Pseudomonas putida*, *Mycobacterium intracellulare*, and the fungus *Fusarium oxysporum* (Supplementary Table 5). The assemblies confirmed that anthropogenically impacted wetland sites harbored more pathogens and in higher relative abundance than natural sites. This might suggest an increased public health risk associated with anthropogenically impacted wetland sites^79^. Among the detected pathogens, *Aeromonas veronii* and *Vibrio cholerae* were most prominent, and might be a concern for public health ^80–82^: At a wetland adjacent to a duck farm, we found high relative abundances of *A. veronii*, an opportunistic pathogen that can cause septicemia in waterfowl and infections in immunocompromised individuals ^83,84^. Similarly, *V. cholerae* emerged as a most prominent pathogen at a wetland site in Spain in proximity to a wastewater treatment plant; here, the high relative abundance might be explained by warm, nutrient-rich conditions ^85^.

We additionally identified the presence of several AMR genes, including clinically relevant ones. Among the 19 detected AMR genes, beta-lactamases were the most abundant ones, and were mostly detected at the wetland site close to duck farms in France: This site was enriched for *bla*_OXA_, *bla*_OXA-956_, and *bla*_FOX_, and also harboured the carbapenemases *cphA* and *cphA1*. The relatively long nanopore sequencing reads (in our metagenomic dataset: median read length of 900 bases) further have the advantage of providing genomic context around the AMR gene from the same physical DNA fragment. This context can be leveraged to assign the gene to its microbial host and therefore assess the potential health consequences of the detected AMR factor. We found that the pathogenic *A. veronii* carried several beta-lactamase genes, including *bla*_OXA_ and *cphA* with known public health relevance ^86,87^ and which are consistent with known *Aeromonas*-associated beta-lactamase repertoires ^87–89^. In cases where the taxonomic resolution does not allow for species-level assignments, additional information can be leveraged to better understand the potential impact of detected AMR genes. For example, we detected the *bla*_VCC-1_ and *bla*_VCC_ genes on sequencing reads assigned to the *Vibrio* genus in the same freshwater samples where other reads pointed to a dominance of pathogenic *V. cholerae*—which might suggest the presence of resistant pathogens at these sites ^90^. In addition, we know that *V. cholerae* is one of the few *Vibrio* species that can survive in freshwater ^91,92^ further corroborating the link between pathogenicity and resistance.

We further confirmed all the described pathogen-resistance associations based on *de novo* assemblies, which provide more robust evidence than individual reads. Read-based analyses, however, detected more AMR genes than assemblies (210 versus 33 genes)—which could be false-positive hits, or they could be a consequence of assembly-based approaches suffering from imperfect retention of AMR genes compared to read-based analysis ^31^. We, however, also found additional resistance genes on the assembly level, namely *nreB* on *Acinetobacter pitii*, and *arsB* and *arsC* on *Acidiphilium multivorum*. The presence of the arsenic resistance genes *arsB* and *arsC* in freshwater sources close to duck farms is interesting since arsenic-based compounds were historically used in agriculture, including as growth-promoting additives in poultry and swine feed ^93,94^. The additional detection of these genes on the assembly level could be explained by the increased sequence identity of the AMR genes on the contig level (*nreB*: 91.2%; *arsB*: 90.1%; *arsC* (mean): 94.3%) in comparison to the read level (*nreB*: 86.6%; *arsB*: 86.7%; *arsC* (mean): 83.0%) where the identities closely missed the default identity threshold by the resistance gene annotation tool ^60^. While this shows that assemblies can improve the accuracy of certain genomic regions, such *de novo* assemblies have to be understood as computational constructs that fuse sequencing data from different microbial cells. When it comes to pathogen associations of AMR genes, read-based resistance detection should therefore remain the ground truth.

We further performed random amplification and rapid transposase-based nanopore sequencing (Rapid SMART-9N) to assess the RNA virome captured by the passive samplers. A diverse group of viral families were detected, with the *Picornaviridae* family as the only one with known mammalian and avian hosts. Picornaviruses are non-enveloped viruses that are environmentally stable and often indicate fecal contamination in surface waters ^95^. Here, *Picornaviridae* were found at all anthropogenically impacted wetland sites where viral contigs were obtained, in contrast to natural sites where they were found in only two out of the six locations. We also detected the *Dicistroviridae* family, which includes many insect-infecting viruses such as bee-associated viruses with relevance for pollinator health ^96^. Several contigs were assigned to the *Nodaviridae* family, which includes fish pathogens (betanodaviruses) associated with viral nervous necrosis in marine and freshwater species ^97^, and to plant-infecting families such as *Tombusviridae* and *Virgaviridae* ^98,99^. The remaining families were mostly bacteriophages or mycoviruses, which are expected in these environmental samples ^100^.

We were not able to reliably classify many viral sequences to species level. More than half of the viral sequences remained completely unclassified, likely reflecting viral dark matter with high levels of unknown viruses ^101^. This taxonomic resolution could potentially be improved in the future by increasing viral coverage, which could also help with the detection of new viral species through approaches such as phylogenetic placement analysis to resolve divergent lineages ^102–104^. To achieve higher viral coverage, sequencing depth could be increased, or ribosomal RNA could be depleted; however, both approaches remain cost-prohibitive at the time of this study.

The AIV-comprising viral family *Orthomyxoviridae* was not detected at any site, probably because AIV concentrations were below the limit of detection of our random-amplification method. As AIV is, however, a pathogen of increasing global significance which is known to be transmitted along major migratory routes through avian hosts ^105^, we additionally applied a targeted approach by qPCR to detect and characterize AIV strains. We found that the torpedo-shaped passive samplers were able to detect the presence of AIV at a third of our sampled wetland sites. We therefore confirmed these samplers as a cost-effective and easy-to-implement tool for avian influenza detection, as previously shown by ^23^. The majority of our AIV detections were from wetland sites in the area of Toulouse, France, and in close proximity to high concentrations of intensive duck farming for foie gras production. One of the duck farms in whose proximity we sampled suffered from highly pathogenic AIV (HPAIV) outbreaks in 2022 ^106^, and we have previously detected recurrent LPAIV cases in that region (data not shown). The close proximity of AIV-carrying migratory bird and their wetland habitats to poultry farming is a known concern for public health, with genetically depauperate domesticated birds contributing to the spread and mutations of pathogens in general ^9^ and to the emergence of HPAIV specifically ^11^. Our preliminary findings together with this potential public health risk might therefore warrant temporally and spatially larger-scale wetland monitoring endeavors using labor- and cost-efficient passive water sampling approaches.

We were only able to subtype one of the detected AIVs by amplification using universal primers and subsequent nanopore sequencing ^44^, probably due to the low viral quantity and high RNA fragmentation in the water samples ^107,108^. This approach could potentially be optimized to obtain whole-genome data of detected AIV in a more robust manner, for example by using more specific primers to avoid untargeted amplification ^109^, applying a multiplex PCR targeting several regions of the HA segment to increase sensitivity ^110^, or using hybridization probe capture for targeted enrichment ^107^.

We next leveraged the same DNA that we extracted for metagenomic analyses to sequence eDNA for potential avian host detection. Using 12S rRNA amplicons and the capability of nanopore sequencing technology to also profile short reads of less than 100bp at high sequencing accuracy ^38,39^, we detected the presence of several bird species. For example, we found evidence of the geese genus (*Anser spp*.) in a wetland close to Ciudad Real, Spain, where we also detected AIV, and where we have additionally recently detected AIV in geese fecal samples ^111^. In a wetland adjacent to duck farms in France, we found dominance of of duck (*Anas platyrhynchos*) eDNA, and phylogenetic analysis of the recovered AIV HA segment pointed to a H4 viral subtype that was closely related to other H4 subtypes previously detected from *Anas platyrhynchos*. While these results might suggest possible host-inferences from non-invasive samples, no direct link can be established given the presence of eDNA of several bird species in most of the samples. In our case, only at one natural wetland site in France, only a single avian taxon, the cormorant genus (*Phalacrocorax* spp.), was detected alongside AIV and could therefore potentially be inferred as the most likely host. Just such a detection of potential hosts can give valuable information, for example on what wildlife is at risk of being infected and of serving as pathogen reservoirs and vectors, or which endangered species could be impacted in their recovery. While it has been suggested that shotgun sequencing of environmental samples can detect the presence of vertebrates without any need for targeted amplification ^38^, we here found that at a standard sequencing depth none of the avian species detected by eDNA were detected by our metagenomic approach, suggesting a need for eDNA approaches for concurrent sensitive host detection.

In conclusion, we demonstrate that passive water sampling followed by nanopore sequencing provides a holistic and cost-efficient approach for real-time surveillance of wetland ecosystems. With a single non-invasive sampling effort, we were able to detect clinically relevant bacterial pathogens and their associated AMR genes, recover an overview of the RNA virome, identify and subtype AIV, and infer potential avian hosts through vertebrate eDNA. In our study of natural and anthropogenically impacted wetland sites across the East Atlantic Flyway, we found that pathogen detection was consistently higher in anthropogenically impacted wetlands, with robust long-read *de novo* assemblies confirming their presence. This integrated approach highlights how passive samplers capture temporally and potentially spatially cumulative signals and, when combined with the versatility of nanopore sequencing, allow to recover long-read genomic contexts and *de novo* assemblies as well as accurate short-read metabarcodes, with potentially valuable insights into the One Health interface between the environment, wildlife, livestock, and human populations.

## Data and code availability

All nanopore sequencing raw data have been made available at ENA (PRJEB96272). The AIV H4 obtained consensus sequence was uploaded to GISAID (“A/environment/Occitania/1/2024”; EPI_ISL_20139219). All code has been made publicly available via Github: https://github.com/ttmgr/Wetland_Health.

## Acknowledgements

We acknowledge the use of LLMs for support in code development. All code was reviewed and verified by the authors.

## Author contributions

T.R, A.P., and L.U. designed the experiment. T.R, A.P. conducted the sampling, processing, and data analysis under L.U.’s supervision. T.R, A.P., and L.U. wrote the manuscript. All authors approved the manuscript.

## Financial Disclosure

This study was funded by a Helmholtz Principal Investigator Grant awarded to LU, and a German One Health Platform Pilot Project grant awarded to AP and LU (2824HS010). Sampling and primary processing of samples at Spanish wetlands was co-funded by projects MCIN/AEI/10.13039/501100011033 (grant no. PID2020-114060RR-C32) and PID2023-149441OR-C33 financed by MICIU/AEI /10.13039/501100011033 and for FEDER, UE

## Supplementary Information

**Supplementary Table 1. Sampling data of all wetland sites**, including site ID, country, environment category, site number, detailed site descriptions, GPS coordinates, and environmental parameters (Methodology). An interactive Google Earth map can be found here: https://www.bit.ly/locations_pathogen.

**Supplementary Table 2. DNA metagenomics metrics of all samples**, including total DNA yield and concentration, total number of sequencing reads, number of sequencing reads after filtering, read length distribution (median and N50), read-based taxonomic classifications, *de novo* assembly statistics using metaFlye and nanoMDBG (number of contigs, number of bases in contigs, median contig length, and N50 contig), Prokka annotations, and contig-based taxonomic classifications (Methodology).

**Supplementary Table 3. RNA virome metrics of all samples**, including RNA concentration and total yield, and cDNA concentration and total yield post-SMART-9N, and total number of sequencing reads, read length distribution (median and N50), viral mapping results using DIAMOND blastx (number of reads and contigs mapping to viral kingdom and families, and percentage of reads mapping to viruses), and assembly statistics using nanoMDBG (number of contigs, number bases in contigs, median contig length, and N50 contig) (Methodology).

**Supplementary Table 4. Pathogen species detection of all samples based on sequencing reads**: sampling site, sample ID, pathogen species, and number of reads mapping to each pathogen (Methodology).

**Supplementary Table 5. Pathogen species detection of all samples based on nanoMDBG assemblies**: sampling site, sample ID, pathogen species, and number of contigs mapping to each pathogen (Methodology).

**Supplementary Figure 1.**
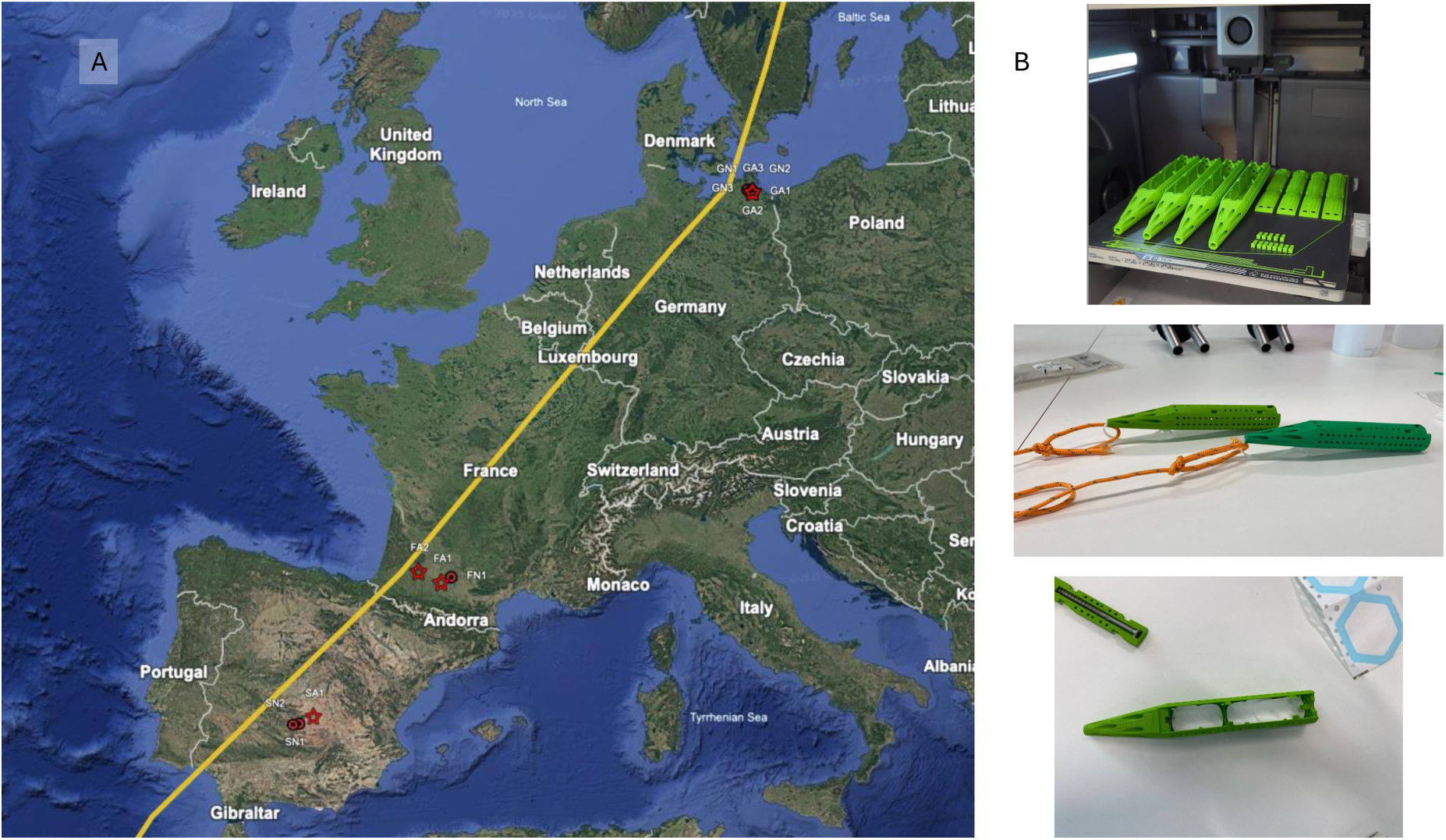
Sampling locations and tools. **(A)** Geographic overview of the twelve sampled wetland sites along the East Atlantic Flyway. Anthropogenically impacted sites are indicated by red stars and natural wetlands by red circles. The main migratory corridor of the East Atlantic Flyway is shown as a yellow line. The interactive Google Earth map can be found here: https://www.bit.ly/locations_pathogen. **(B)** Photos of the 3D-printed torpedo-shaped passive water samplers.

**Supplementary Figure 2.**
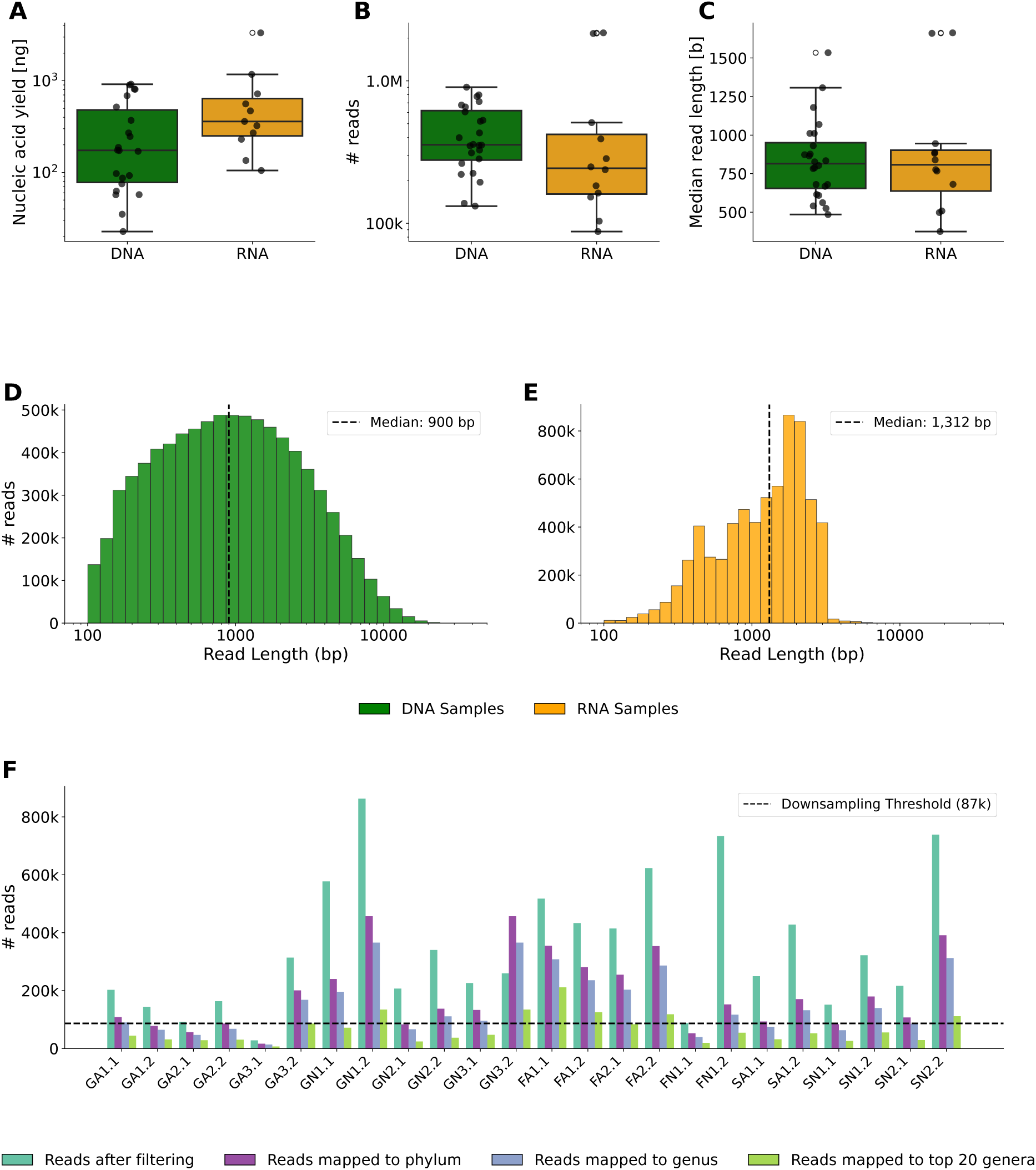
Quality control summary of DNA and RNA sequencing data. The boxplots compare **(A)** nucleic acid yield, **(B)** total raw reads, and **(C)** median read length between DNA- and RNA-based sequencing libraries. Histograms show the sequencing read length distribution for all **(D)** DNA, and **(E)** cDNA reads, with dashed vertical lines indicating the median read lengths. **(F)** Number of total metagenomic reads after filtering and subset of reads mapping to the phylum and genus levels and to the top 20 genera. The horizontal dashed line indicates the downsampling threshold of 87,000 reads. GA3.1 was excluded for subequent analyses since it did not reach the downsampling threshold.

**Supplementary Figure 3.**
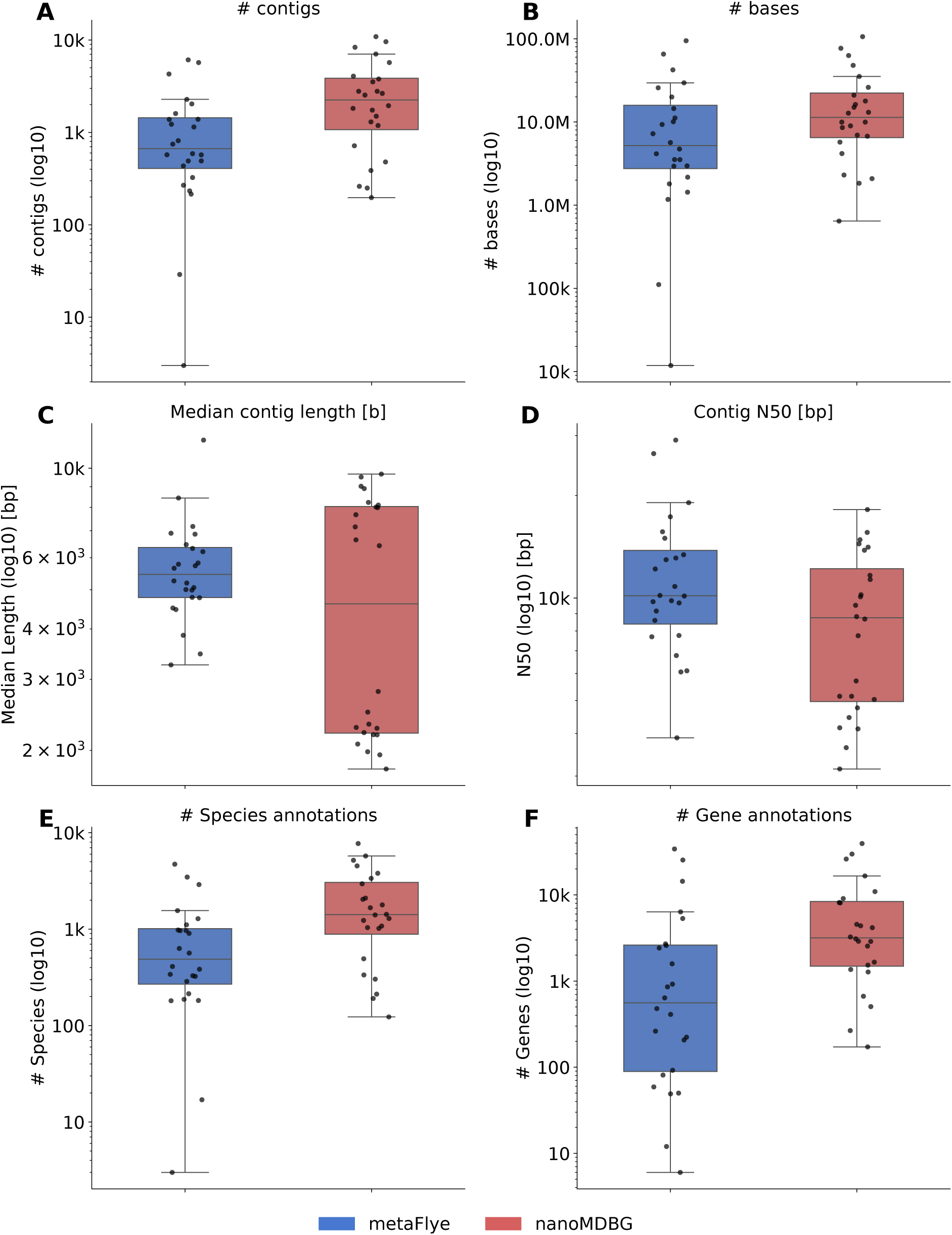
Comparison of *de novo* metagenome assemblers. Comparison of metaFlye and nanoMDBG long-read assemblers across all samples in terms of **(A)** number of assembled contigs, **(B)** number of assembled bases, **(C)** median contig length, **(D)** contig N50 value, **(E)** number of unique detected species, and **(F)** number of gene annotations.

**Supplementary Figure 4.**
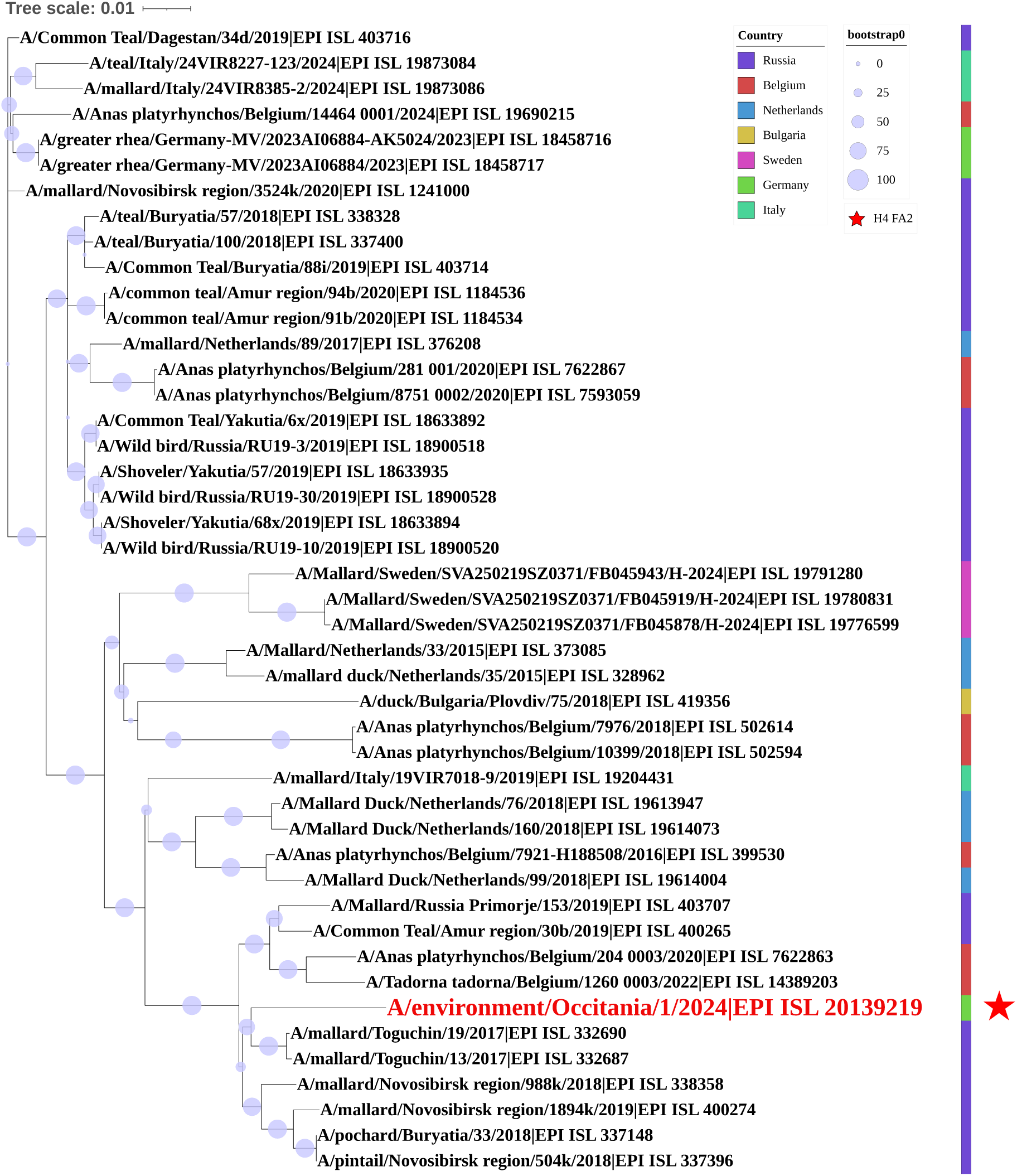
AIV phylogenetic tree. Phylogenetic analysis of the AIV HA H4 segments from our sample FA2 (highlighted in red) with H4 segments from the GISAID dataset in Europe from January 2015 to August 2025. A branch length of 0.01 corresponds to ∼1 substitution per 100 sites. The colour scale represents the country where the viruses were collected.

## Notes

### Competing Interest Statement

The authors have declared no competing interest.

