## Supplementary figures and images for "Real-time genomic pathogen, resistance, and host range characterization from passive water sampling of wetland ecosystems"

### Supplementary_Figure1.pdf

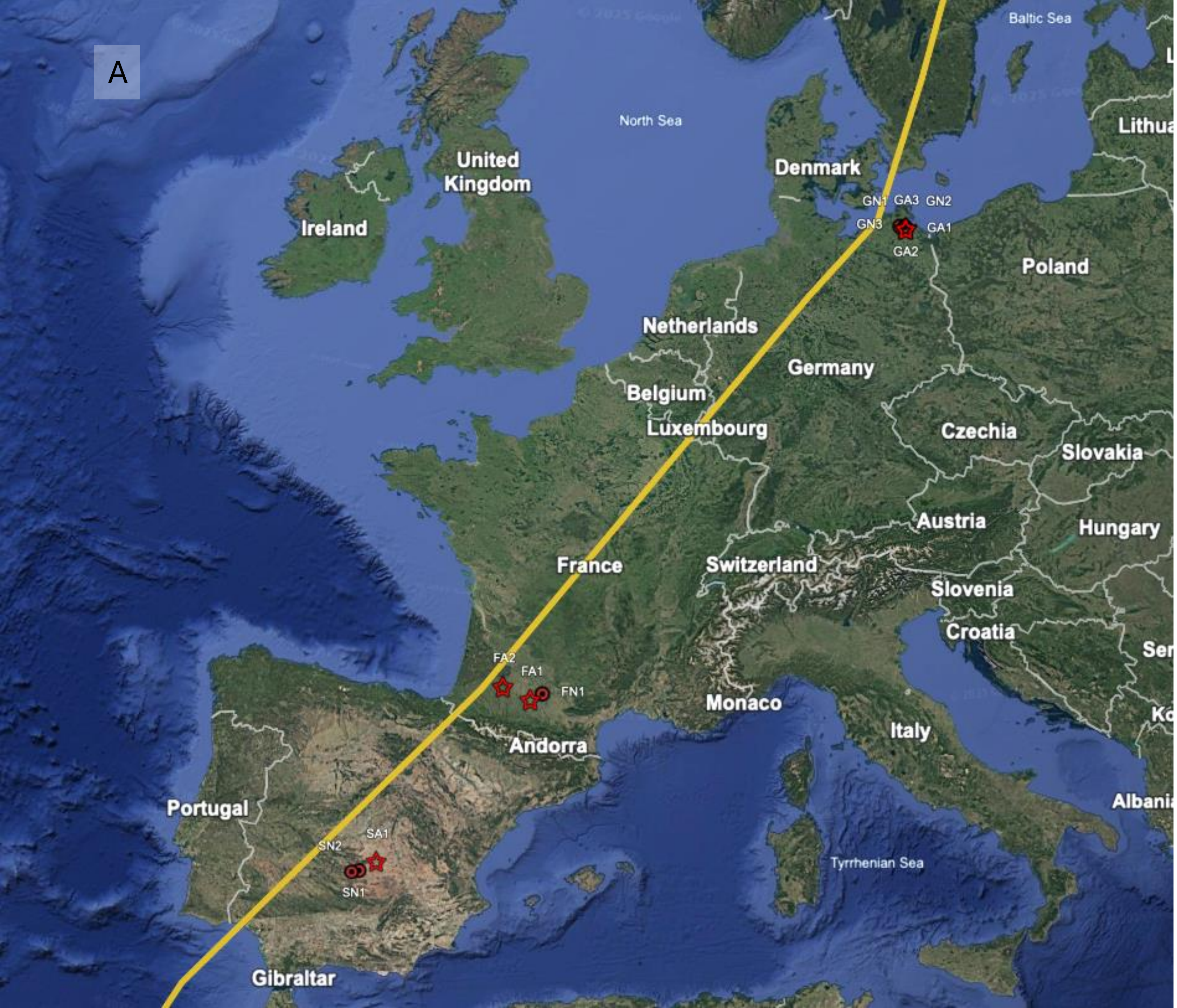

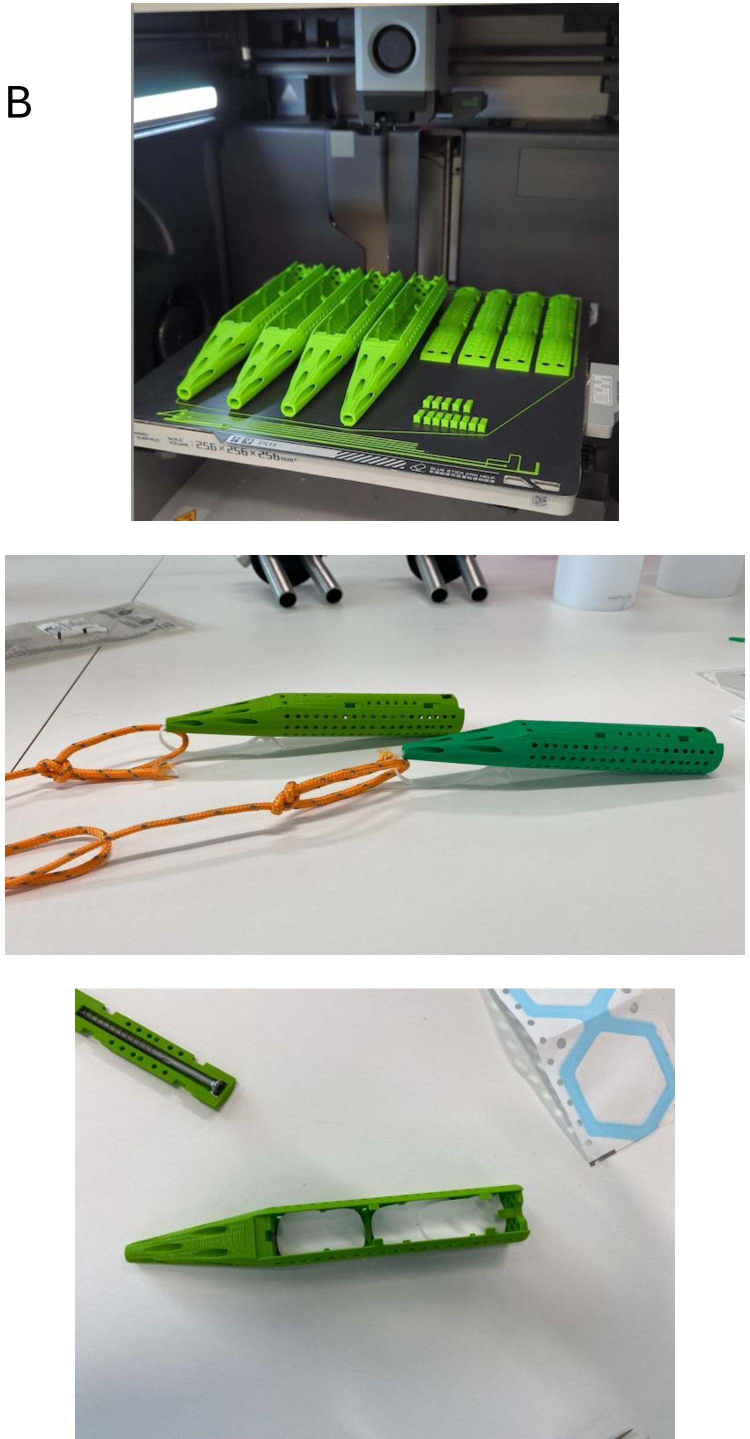

### Supplementary_Figure2.pdf

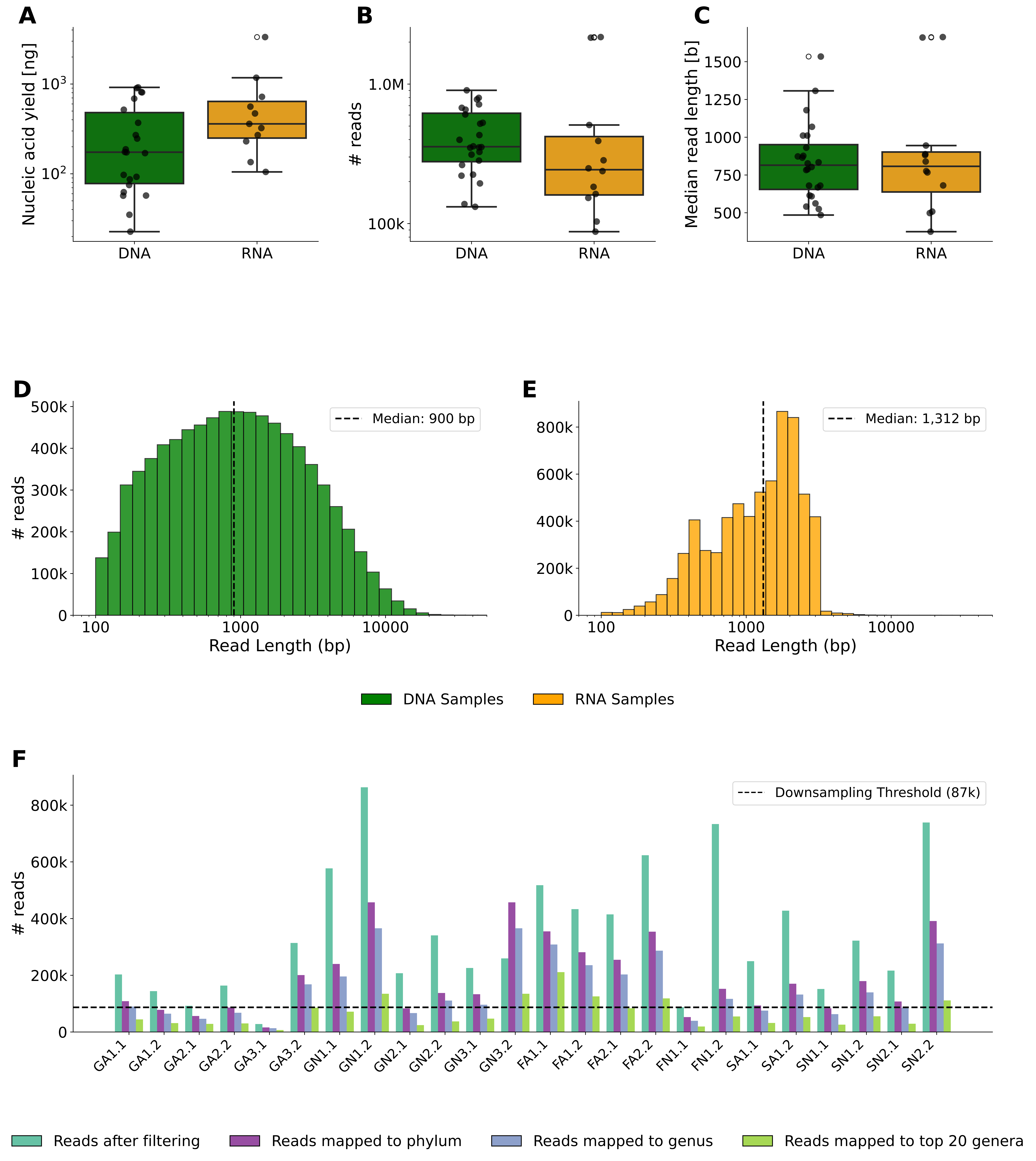

### Supplementary_Figure3.pdf

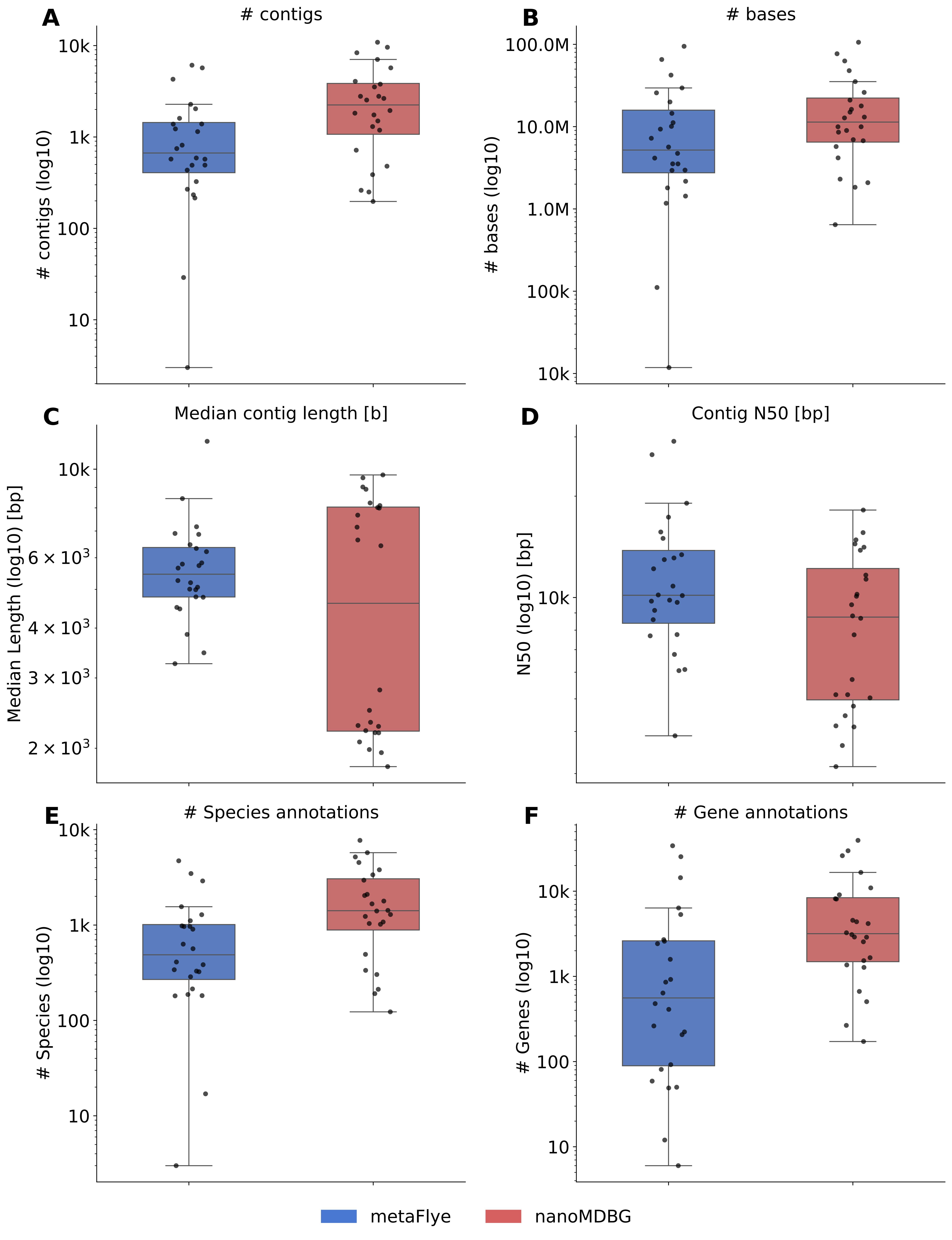

### Supplementary_Figure4.pdf

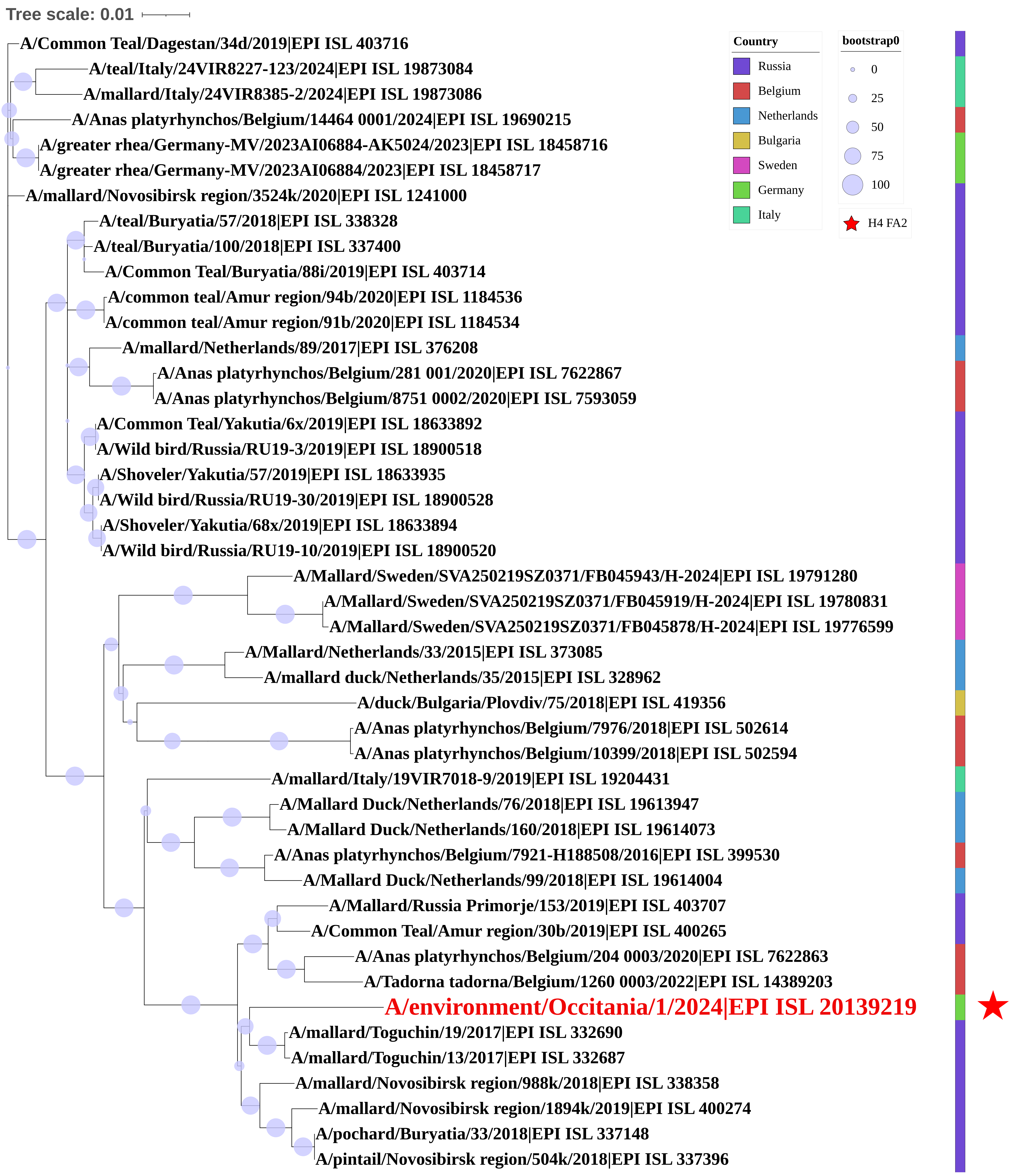
